# In form for a swarm: programmable neutrophil swarming impacts infection outcome

**DOI:** 10.64898/2025.12.03.692046

**Authors:** Salik M. Borbora, Giulia Rinaldi, Tuhin Samanta, Can Cui, Echo Yongqi Luo, Ivanna Williantarra, Debsuvra Ghosh, Holly M. Craven, Nir S. Gov, Milka Sarris

**Affiliations:** Department of Physiology, Development and Neuroscience, University of Cambridge, Downing Site, Cambridge, CB2 3DY, UK; Department of Chemical and Biological Physics, Weizmann Institute of Science, Rehovot, Israel; Roving Researcher Programme, School of Biological Sciences, University of Cambridge, United Kingdom

**Author notes:** These authors contributed equally (Salik M. Borbora, Giulia Rinaldi).

## Abstract

Many migrating cells pattern signalling molecules to support their accumulation in target tissues. During inflammation, neutrophils achieve rapid swarming at sites of injury or infection by generating spatiotemporal gradients of chemoattractants. Whether these self-generated behaviours can be re-programmed to change disease course is unclear. Here, we show that neutrophil chemoattractant patterning machinery is subject to transcriptional reprogramming after microbial experience, leading to enhanced swarming and control of infection at subsequent wound sites in zebrafish. Through in vitro assays, we demonstrate that “trained swarming phenotypes” are cell-intrinsic and independent of the nature of the microbial target. Furthermore, genetic enhancement of 5-lipoxygenase in neutrophils is sufficient to improve neutrophil swarming and resistance to wound infection. Through mathematical modelling, we demonstrate that neutrophil-intrinsic alterations in chemoattractant release are sufficient to recapitulate trained swarming in infection-experienced animals. Together, these data suggest new routes for reprogramming cell migration in disease settings, via manipulating their ability to shape chemoattractant landscapes.

## Introduction

Directed cell migration is essential for tissue development and surveillance. Guidance of migrating cells relies on interpretation of gradients of chemoattractants within tissues. Traditional mechanistic models portray migrating cells as receivers of information, responding to chemoattractant landscapes patterned by extrinsic tissue cells. However, several lines of evidence highlight an important role for self-organisation in the process, through cell-autonomous generation of chemical gradients by migratory cells^1–5^. For example, during development, migrating tissue collectives sculpt chemical gradients through asymmetric distribution of dedicated chemoattractant scavenging receptors^3,5^. In cancer metastasis, melanoma cells disperse via catabolism of ligand and local micro-gradient generation^6^. In chemotactic mazes, amoeboid cells detect dead ends using breakdown of chemoattractant^7^. During immune responses, leukocytes produce autologous chemical gradients through chemoattractant scavenging or through autocrine/paracrine secretion. Finally, modification of ligands by migrating cells can generate complex patterns such as reverse chemotaxis^13^. A fundamental question raised by these observations is whether cell guidance is autonomously programmable, in a manner sufficient to impact disease at the tissue and organism scale.

To explore this question, we focus on the paradigm of neutrophil swarming, as a self-organised migration behaviour. Neutrophils are first responders to tissue injury and infection and directly fight microbes through a variety of effector mechanisms, such as phagocytosis, secretion of antimicrobial molecules, and release of neutrophil extracellular traps (NETs)^14^. Neutrophils also produce chemokines and cytokines that influence the accumulation and function of other cells and ultimately shape the course of inflammation and disease. The recruitment of neutrophils is initially prompted by primary signals produced by tissue cells afflicted by injury or infection^14^. Thereafter, neutrophils begin to secrete further chemoattractants, generating tissue-scale gradients and/or waves of chemoattractants, which amplify the range of recruitment and determine the final magnitude of the response^9–12,15–17^. The process of neutrophil swarming is highly conserved and thought to impart functional benefits in fighting infection, particularly in the face of large and focal targets, such as wound infection or fungal invasions^10,18,19^. Specifically, swarms are characterized by rapid kinetics of recruitment and dense accumulation of neutrophils^9–12,15^, which in principle would support prompt containment of pathogens, physical insulation of the pathogen-afflicted area and local concentration of antimicrobial molecules. On the other hand, neutrophil accumulation is also known to prolong pathological inflammation when not properly regulated and resolved^14,20^. Therefore, the mechanisms and rules of organisation underlying neutrophil swarm development have important functional implications in disease.

Several conserved mechanisms drive the initiation and evolution of neutrophil swarming. This includes the key neutrophil-derived chemoattractants driving swarm development (notably, the eicosanoid leukotriene B4 (LTB4) and the chemokine Cxcl8^9,10,15,21^), signal inducers of chemoattractant secretion (such as coordinated calcium signals^10,11,22^) as well as factors that self-limit the evolution of swarms (such as receptor desensitization, lipoxins, resolvins and NADPH activity^11,15,23,24^). Additional contributing factors have also been described, such as damage associated molecular patterns (for example ATP and fMLP^10,12^), complement products^25^, mechanical signals^26,27^, transcellular synthesis^28^, neutrophil NETosis and target cloaking by other immune cells^21,29^. Despite the overall conservation of neutrophil swarming and associated mechanisms, the dynamics of neutrophil swarm behaviours within a given experimental system show significant heterogeneity, ranging from small, short-lived clusters, to persistent and larger clusters^10,30^. Although some of this heterogeneity can be attributed to variation in the intensity and composition of tissue-derived signals^23,29^, it remains unclear whether cell-intrinsic properties account for variation in neutrophil propensity to swarm and whether these are subject to re-programming.

Immune cell gene expression and function is re-programmed naturally through immune experience but also therapeutically through genetic engineering. In the first case immune cells undergo ‘training’, by metabolic rewiring and epigenetic marks that poise their gene expression profile and responsiveness to a subsequent insult^31–33^. Neutrophils have been shown to undergo such durable training after exposure to primary infection, through epigenetic re-programming in relevant long-lived progenitors^34,34–37^. Innate immune training increases resistance to infection in a variety of tumour and infection settings and involves a range of epigenome and transcriptome changes^34,35,38,39^. Whilst it is well-established that neutrophils enhance their killing capacity and associated gene signatures after immune training^34,40–42^, it remains unclear to what extent their ability to swarm can also undergo re-programming under these conditions. On the other hand, re-programming immune function through genetic engineering is exemplified in tumor immunotherapies. Specifically, chimeric antigen receptors (CAR) can be engineered in immune cells to programme their ability to recognise and kill certain tumours^43^.

Whilst this has been developed to the largest extent for T lymphocytes^43^, new approaches for CAR therapies are geared towards the manipulation of myeloid cells, including macrophages and neutrophils^44–47^. However, the ability to genetically reprogramme the infiltration of immune cells to target diseased tissues remains a challenging frontier. In this context, the manipulation of self-generated gradients is a relatively unexplored but interesting route of manipulation, as it offers the potential to modify the composition of inflammatory niches through the reprogramming of ligands delivered by immune cells to the site.

Here we exploit the zebrafish as a tractable model to explore the link between neutrophil behaviours at the microscopic level with disease outcome at the whole-organism scale. We show that training challenge of zebrafish with non-pathogenic microbes enhances resistance to subsequent wound infections and that, under these conditions, neutrophils exhibit enhanced swarming prowess. These trained swarming phenotypes are cell-autonomous and associated with transcriptional upregulation and epigenetic marking of swarm-related genes (LTB4 biosynthesis pathway). Through genetic engineering and mathematical modelling, we show that changes in chemoattractant release are sufficient to enhance neutrophil swarming and resistance to wound infection.

## Results

### Training through microbial challenge alters expression of swarm machinery and enhances resistance to wound infections

To investigate the dynamics and functional consequences of neutrophil swarms, we employed a wound infection model. *Pseudomonas aeruginosa* is an opportunistic bacterial pathogen, present in water and soil, able to colonise wounds in multiple species. In humans, wound infections can occur through contact of injured tissue with contaminated water or surfaces^48,49^. These infections can be severe and lead to sepsis and mortality, particularly in immunocompromised individuals or in the case of large surgery-associated infections^48,49^. We previously introduced a model whereby zebrafish larvae are subjected to wounding in the presence of aqueous medium contaminated with *P. aeruginosa* PAO1 strain^10^. This experimental model is advantageous in recapitulating a natural route of infection, in comparison to common microinjection routes^50^, and provides a relevant setting to study the role of neutrophil swarming in defence.

To induce innate immune training we injected non-pathogenic, CFP-expressing *Escherichia coli* MG1655^51^ in the otic vesicle at 2 days post-fertilisation **(Fig. 1A)**. We found this pre-challenge increased the resistance to wound infection by *P. aeruginosa*, two days after the first challenge, as compared with controlled PBS-pre-challenged larvae **(Fig. 1B-D)**. This was reflected both in terms of disease symptom severity (see methods section for scoring system) and survival **(Fig. 1B-D)**. Notably, in the absence of wounding, larvae were not susceptible to infection in either PBS *or E. coli-*pre-challenge, confirming the opportunistic nature of infection **(Fig. S1A)**. Varying the dose of *E. coli* injection showed that a minimum dose was required to trigger protective effects **(Fig. S1B)**. One way of distinguishing innate immune ‘training’ from immune ‘priming’ is by assessing whether the protective effects persist after the immune response to the first challenge has passed its peak or resolved. To explore the nature of the protection observed, we determined the resolution kinetics of the immune response after the first challenge, by measuring titres of *E. coli* and levels of inflammatory cytokines **(Fig 1E)**. Both markers indicated that the response to *E. coli* had largely subsided by the time the second challenge was given **(Fig 1F-J)**.

**Figure 1:**
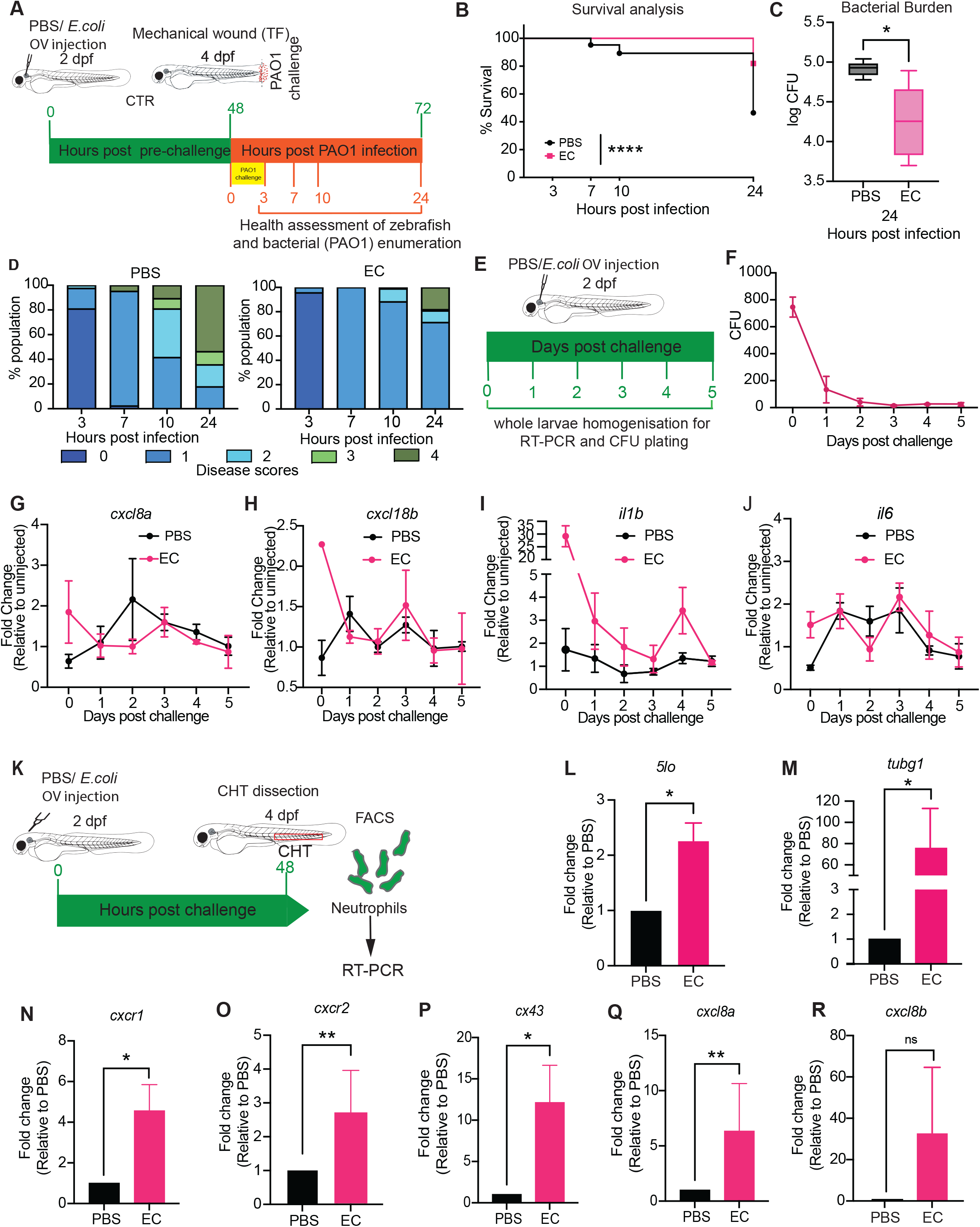
Immune training enhances resistance to wound infections and reprograms genes associated to swarming. A. Schematic of the training experiment. Zebrafish larvae pre-challenged by PBS or a dose of *E. coli* in the otic vesicle (OV) at 2 days post fertilization (dpf) stage and subsequently subjected to mechanical wounding in the tail fin (TF) and *P. aeruginosa* PAO1 infection at 4 dpf. B. Survival curves from control (PBS) and trained (EC) zebrafish larvae upon *P*. *aeruginosa* infection following mechanical wounding (MW). (Representative of three independent experiments with n ≥ 36 larvae per group in each experiment). *Log-rank (Mantel-Cox) test; ****P < 0.0001*. C. Bacterial burden of *P. aeruginosa* in the control and trained zebrafish larvae at 24 hpi (hours post infection). (Representative of three independent experiments, with n ≥ 9 larvae per group used for CFU analysis in each experiment. (mean ± SD). CFU, colony forming units. *One-way ANOVA with Tukey’s multiple comparisons test, *P < 0.05* D. Disease progression in the control and trained zebrafish larvae at the indicated time points. Representative of three independent experiments with n ≥ 36 larvae per group in each experiment. E. Schematic detailing the assessment of *E. coli* burden and cytokine profile in whole zebrafish larvae after primary challenge (PBS/ *E. coli*) for trained immunity induction. F. Bacterial counts at the indicated time points after injection of *E. coli* in the otic vesicle (OV) of the zebrafish larvae. Representative of three independent experiments, with n ≥ 12 larvae for each time point. G-J. Change in the expression profile of the indicated cytokines in naive and trained zebrafish larvae at the specific time points after primary challenge. Representative of three independent experiments, with n ≥ 8 larvae at each time point. K. Schematic describing the isolation of neutrophils by Fluorescence Activated Cell Sorting (FACS) from the hematopoietic site (CHT: Caudal hematopoietic tissue) of zebrafish larvae, followed by transcript analysis using RT-PCR. L-R. Change in the expression profile of the indicated genes in naive and trained zebrafish larvae at 4dpf (2 days after primary challenge). Quantitative PCR data are presented as mean ± S.E.M., with expression levels normalised to PBS-treated control groups (n=5 independent experiments, with each replicate representing a pool of 150-200 larvae at 4 dpf). Statistical significance was assessed using the Mann-Whitney test/ t-test (*p < 0.05, **p < 0.01).

Neutrophil training is characterised by changes in transcriptomic and epigenetic signatures in differentiated neutrophils and progenitors^37^. Since the training challenge led to resistance of infection in a heterologous tissue site (tail versus ear), we hypothesised neutrophils from hematopoietic sites might be centrally altered. To explore this, we isolated neutrophils from the caudal hematopoietic tissue (CHT), principal site of hematopoiesis of the fish at this stage **(Fig. 1K)** and found that CHT-derived neutrophils showed increased expression of genes associated with inflammation and cell migration, in particular *cxcl8a*, *5-lo* and cytoskeletal proteins, such as *tubg1* **(Fig 1L-R)**. This suggested that neutrophils may be centrally altered by training in a manner that changes their gene expression profile. To explore epigenetic changes in genes associated to swarming, we mined published datasets from neutrophil training by *S. flexneri,* which is highly related to *E. coli* phylogenetically^52^ ^42^. Zebrafish neutrophils undergo epigenetic reprogramming upon sub-lethal *Shigella flexneri* infection, and this is associated with enhanced resistance to subsequent *S. flexneri* infection. We found that H3K4me3 marks were induced in *Shigella*-trained animals in gene loci associated to eicosanoid (LTB4) synthesis and production, namely *5lo/alox-5*, *lta4h* and *cyp4f3 (***Fig S1C**).

Together, these analyses revealed that innate immune responses to wound infections can be trained by prior microbial challenge and suggested that, among the many pathways of neutrophil function, genes associated with cell migration and swarming can be subject to reprogramming.

### Neutrophil swarming is enhanced through prior training challenge and contributes to disease resistance

The changes in gene expression patterns in neutrophils upon training suggested alterations in migratory behaviour. To explore this, we exploited a laser wound model^10^, in which imaging of wound infections in the early time periods of infection (0-2 hours post infection-hpi) can be coupled to monitoring of disease over a longer timescale (up to 24 hpi) in the same larvae **(Fig. 2A)**. We compared the behaviour of neutrophils upon wounding and found that in PBS-pre-challenged animals, neutrophils exhibited both small-scale clustering and large-scale clustering **(Fig. 2B**, examples 1 and 2 for small and large scale, **Movie S1)**. By contrast, in *E. coli*-pre-challenged animals, we observed consistent large-scale clustering **(Fig. 2B**, examples 1 and 2 for large scale clustering, **Movie S1)**. To understand differences in migratory behaviour, we plotted the average neutrophil radial speed **(Fig. 2C)** and speed **(Fig. 2D)** over time in PBS versus *E. coli-*pre-challenged fish. *E. coli-*pre-challenged fish showed enhanced migration speeds towards the wound throughout the first 2h after wounding **(Fig. 2C-D).** These differences were not explained by different number of neutrophils in the tissue before wounding (**Fig. 2E**). In relation to infection outcome, we found lower bacterial burden in PBS-controls in these laser-wounded larvae (**Fig. 2F**) compared to mechanically-wounded larvae (**Fig 1C**), which could be due to the smaller size of laser wounds. Nevertheless, we confirmed in this laser-wound assay, PBS pre-challenged, control fish were more susceptible to PAO1 infection than *E. coli* pre-challenged fish **(Fig. 2F).**

**Figure 2:**
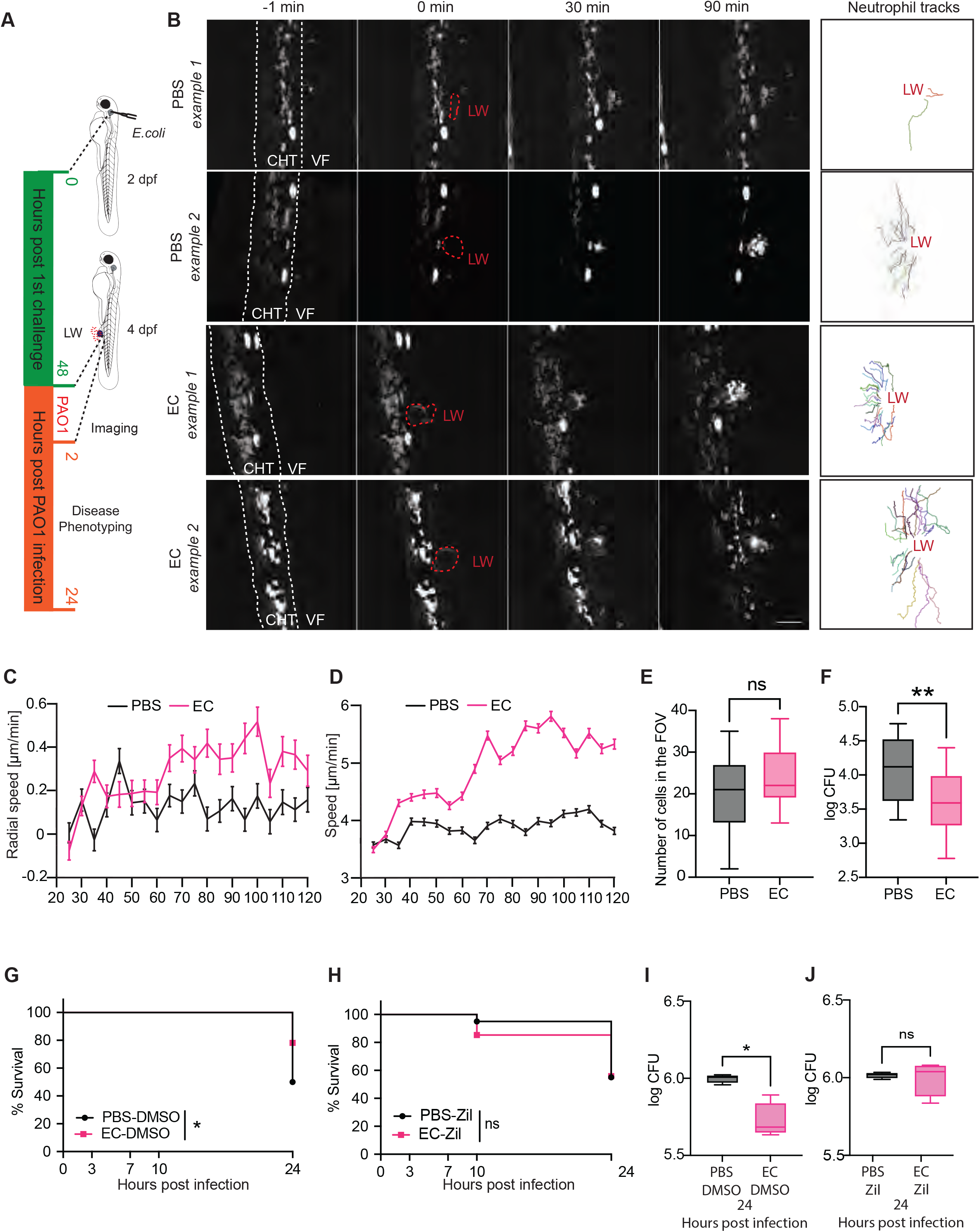
Immune training induces alterations in neutrophil swarming that are beneficial for bacterial defence. A. Schematic of experiment with control, PBS pre-challenged or trained, E. *coli* pre- challenged (EC) zebrafish larvae subjected to laser wounding (LW), followed by infection with *P. aeruginosa* (PAO1) and simultaneous confocal imaging. B. Time lapse images of representative movies of control and trained larvae at 1 min prior LW (-1 min), at the time of LW (0 min), at 30 min and 90 min post LW. White dashed lines correspond to the CHT: caudal hematopoietic tissue and VF: ventral fin. Red dashed lines correspond to the site of the LW. On the far-right, neutrophil trajectories of same movie are depicted wherein each colour corresponds to a different neutrophil trajectory. Scale bar 50 μm. C. Radial speed quantification of tracked cells in control (PBS) (black) and trained EC animals (pink) overtime after laser wound and infection. (PBS, n=16) (black) and trained EC (n=16) animals (pink). D. Speed quantification of tracked cells in control (PBS, n=16) (black) and trained EC (n=16) animals (pink) overtime after laser wound and infection. E. Initial number of neutrophils in each zebrafish larvae in the FOV: field of view before laser wounding across both PBS pre-challenged and *E. coli* pre-challenged larvae that were subjected to laser wounding and infection with simultaneous imaging. n=16 (PBS) and n=16 (EC). Unpaired t test with Welch’s correction. *ns p > 0.05*. F. Bacterial burden of PAO1 in PBS pre-challenged and *E. coli* pre-challenged larvae, 24 hours post infection (hpi), and subjected to laser wounding and infection with simultaneous imaging. n=16 (PBS) and n=16 (EC). *Unpaired t test with Welch’s correction. ** p < 0.01*. G-H. Survival curves from control (PBS) and trained (EC) zebrafish larvae upon PAO1 infection following mechanical wound. Larvae were pre-treated with DMSO or 20µM Zileuton (Zil), 1h prior to wounding and kept in this medium until the end of the experiment Representative of three independent experiments with n ≥28 larvae per group for each experiment in E, and n ≥ 20 larvae in F. Log-rank (Mantel-Cox) test; * p < 0.05, ns p > 0.05. I-J. Bacterial burden of *P. aeruginosa* in larvae from groups in G and H respectively. Representative of three independent experiments with n ≥ 9 larvae per group used for CFU analysis in each experiment. (mean ± SD). CFU, colony forming units. Unpaired t test with Welch’s correction. * p < 0.05, ns p > 0.05.Error bars represent standard error of the mean.

These analyses suggested that *E. coli* training maximises the swarm potential of neutrophils. Our transcriptional and epigenetic analyses revealed variations in 5-LO expression, as a possible mechanism **(Fig. 1L-R** **and Fig. S1C)**. To investigate whether 5-LO activity contributes to wound infection resistance, we used a specific inhibitor for 5-lipoxygenese (5- LO), zileuton, to target the production of the swarm-driving chemoattractant, LTB4. When fish were wounded and infected in the presence of zileuton, the *E. coli*-pre-challenged fish did not show as marked an increase of infection resistance (over PBS-pre-challenged fish), in comparison to when control vehicle (DMSO) was used **(Fig. 2G-J** **and Fig. S2A)**. These differences are compatible with a role for swarm dynamics in infection resistance.

Together, these results indicated that innate immune training leads to changes in neutrophil swarm patterns upon secondary pathogenic challenge, which in turn support infection clearance.

### 5-lipoxygenase enhancement in neutrophils is sufficient to enhance neutrophil swarming and resistance from wound infection

To determine whether reprogramming of neutrophil swarming is sufficient to influence wound infections, we used genetic engineering. We hypothesised that neutrophils expressing maximal levels of 5-LO would have enhanced swarming and infection containment. We first compared the resistance to mechanical wound infection in transgenic larvae in which neutrophils overexpress 5-LO (‘5-LO-Neu’; for 5-lipoxygenase overexpressing neutrophils) versus control wild type larvae. We found that 5-LO-Neu larvae showed enhanced resistance to mechanical wound infection, as measured by an improvement in survival rates, disease score and bacterial burden **(Fig. 3A-C)**.

**Figure 3:**
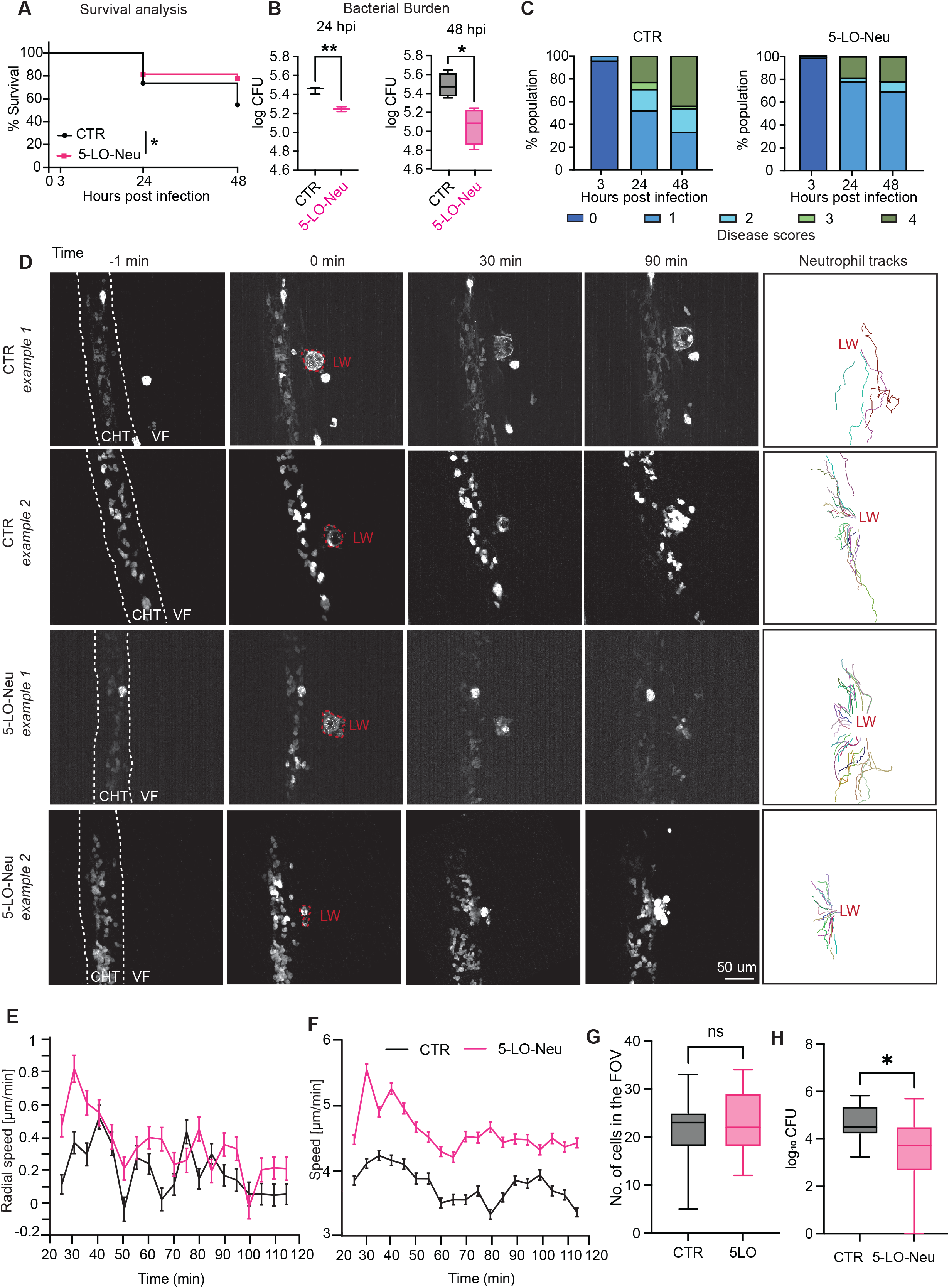
Genetic enhancement of neutrophil LTB4 synthesis is sufficient to alter neutrophil swarm kinetics and infection outcome. A. Survival curves from control (CTR) zebrafish larvae and larvae overexpressing 5-LO in neutrophils (5-LO-Neu) upon *P*. *aeruginosa* infection following MW. Representative of three independent experiments with n ≥ 50 larvae per group in each experiment. *Log-rank (Mantel-Cox) test;* * *p < 0.05*. B. Bacterial burden of fish in survival experiments (A) at the indicated time points. Representative of three independent experiments with n ≥ 9 larvae per group used for CFU analysis in each experiment. *Unpaired t test with Welch’s correction; * p < 0.05, ** p < 0.01*. C. Disease progression in the specified group of larvae at the indicated time points. Representative of three independent experiments with n ≥ 50 larvae per group in each experiment. D. Time lapse images of representative movies of CTR and 5-LO-Neu at 1 min prior to LW (-1 min), at the time of LW (0 min), at 30 min and 90 min after LW. White dashed lines indicated the CHT: caudal hematopoietic tissue and VF: ventral fin. Red dashed line corresponds to the site of the laser wound (LW). On the right column neutrophil trajectories of same movie are represented wherein each colour indicates a different neutrophil trajectory. Scale bar 50 μm. E. Radial speed quantification of tracked cells in control (CTR n=15) (*black*) and 5-LO- Neu (n=17) animals *(pink)* overtime after laser wound and infection. F. Speed quantification of tracked cells in control (CTR n=15) (*black*) and 5-LO-Neu (n=17) animals *(pink)* overtime after laser wound and infection. G. Initial number of neutrophils in each zebrafish larvae in the field of view (FOV) before laser wounding across both CTR and 5-LO-Neu that were subjected to laser wounding and infection with simultaneous imaging. n=15 for CTR and n=17 for 5-LO-Neu. *Unpaired t test with Welch’s correction. ns. p > 0.05*. H. Bacterial burden of PAO1 in larvae from CTR and 5-LO-Neu, 24 hours post infection (hpi), that were subjected to laser wounding and infection with simultaneous imaging. *Unpaired t test with Welch’s correction*. * p < 0.05. n=15 (CTR) and n=17 (5-LO-Neu) Error bars represent standard error of the mean.

To relate neutrophil swarm patterns to disease susceptibility, we combined again laser wounding and live imaging with subsequent monitoring of disease. By observation of the movies, we noted heterogeneity in swarm magnitude across control larvae, which appeared to be less pronounced in 5-LO-Neu larvae **(Fig. 3D**, examples 1 and 2 and **Movie S2**). To quantify these differences, we measured the average neutrophil radial speed **(Fig. 3E)** and speed **(Fig. 3F)** over time in 5-LO-Neu. Neutrophils in 5-LO-Neu larvae were faster, particularly in the first 1h of observation. These differences were not attributed to differential number of neutrophils, as the number of neutrophils in the field of view was comparable across conditions (**Fig. 3G**). As with mechanical wounding, we found that the 5-LO-Neu larvae showed greater resistance to infection in laser wound infection **(Fig. 3H)**. Together these results suggested that 5-LO enhancement in neutrophils is sufficient to enhance neutrophil swarming rates and that this can provide a beneficial outcome in a subset of larvae during pathogenic infections.

Excess neutrophil recruitment can be detrimental for resolution of inflammation and tissue homeostasis. We therefore interrogated whether enhanced neutrophil 5-LO expression could compromise tissue repair and regeneration. To test this, we measured tissue regeneration over 2 days post sterile wound injury. We found that 5-LO enhanced animals did not have impaired regeneration and instead showed a trend for improved regeneration (**Fig. S2B-D**). Thus, the reprogramming of 5-LO in neutrophils did not induce obvious side effects in wound responses.

#### Mathematical modelling predicts cell-intrinsic programming of swarming and bacterial clearance

To quantitatively understand how cell-intrinsic changes in swarm machinery affect swarming, we employed mathematical modelling. Previous studies indicate at least two levels of chemoattractant patterning for swarming^10,11^. Firstly, neutrophils stimulate each other to secrete attractant at the swarm cluster (core production)^10^. Secondly, neutrophils can emit waves of chemoattractant in a relay fashion (active relay)^11,12^. Mathematical models have been previously used to describe the dynamics of signal wave propagation across neutrophils^11^. However, the sufficiency of either core production or active relay to generate neutrophil clustering has not been explored. We hypothesized that core production may be sufficient to generate a primary level of neutrophil clustering, as in our experimental model we observe LTB4-dependent neutrophil clustering and intra-cluster calcium signals, without repetitive active-relay-like calcium waves^10^. This remains compatible with active relays as a further scaling mechanism under different conditions and systems^11,12^.

We modelled the dynamics of swarming using the simplest core production model (for the model equations see Materials and Methods section). Here, the chemotaxis of each neutrophil is described to arise due to the difference in chemoattractant concentration on opposite sides of the cell^53^, with respect to the source at the wound (**Fig. 4A**). We assume the chemotaxis be dominated by a persistent chemoattractant signal that is induced by neutrophils at the wound site, in addition to primary tissue damage signals. We found that this core production by neutrophils at the wound is sufficient to induce clustering of neutrophils at the wound (**Fig. 4B- C, Movie S3**). By contrast, persistent attractant secretion by cells outside the wound led to detrimental clustering of cells outside the wound (**Fig.S3A and B**). This suggested that active relay requires additional assumptions (e.g. periodic fluxes of secretion) to recapitulate swarm kinetics. We therefore focused on the simple core production model, as this was sufficient to reproduce the basic pattern of kinetics of neutrophil swarming in our system (**Fig 4B-C**). In this dynamic system, we could observe both how the bacteria and neutrophil numbers scale over time in a linked fashion (**Fig. 4D**).

**Figure 4.**
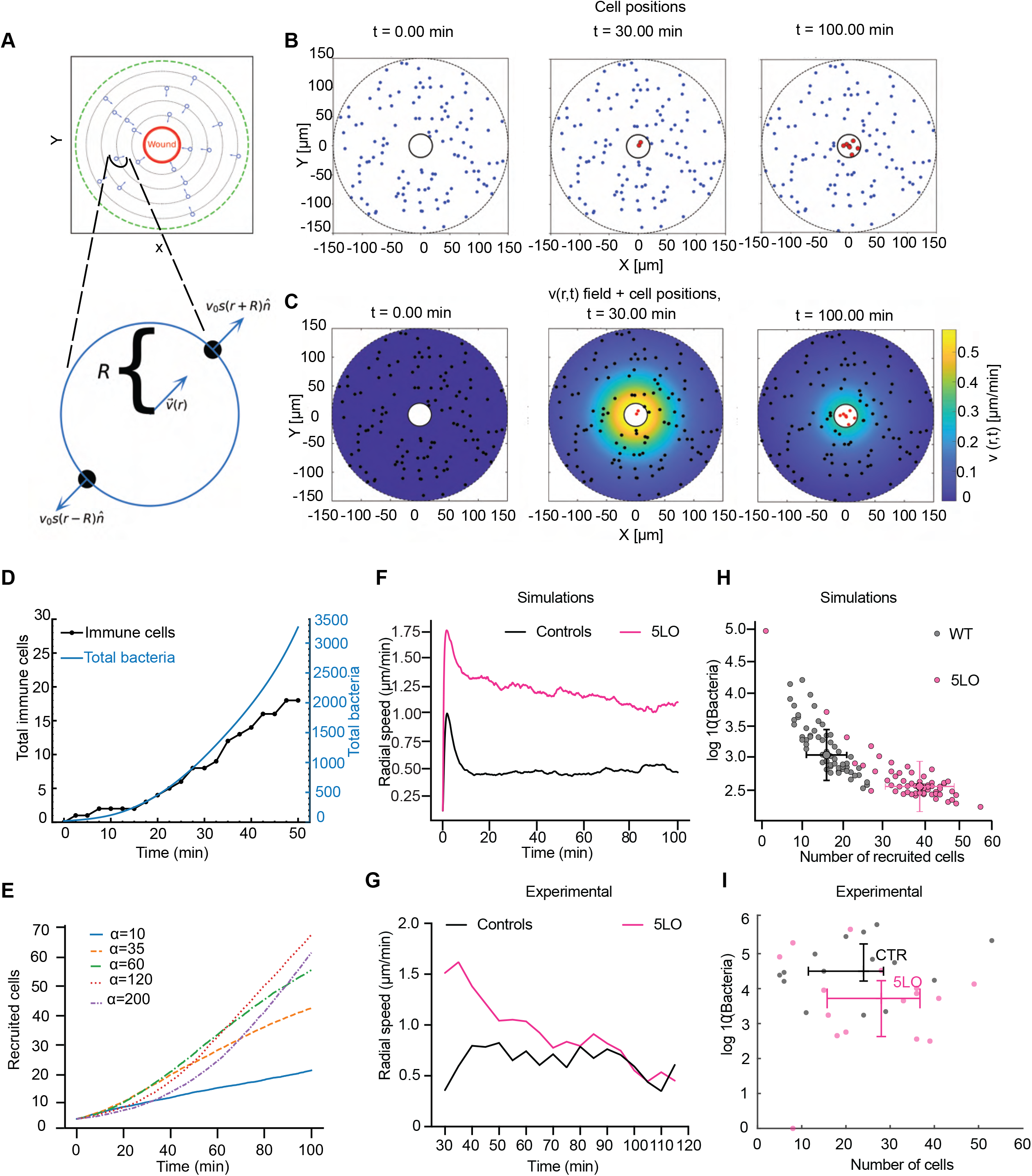
: **Mathematical modelling predicts how variation in neutrophil chemoattractant secretion influences bacterial clearance.** A. Schematic of the simulated domain around the wound. Cells are initially placed randomly with uniform average density, and have a net migration speed towards the wound given by Eqs.(1,2). The simulated domain is bounded at its centre by the wound (red circle), where cells accumulate. At the outer boundary (green dashed line), there is an influx of cells, at random places around the edge, to maintain the density at the edge constant on average. The bottom panel explains how the calculation of the average chemotactic cell speed by the difference between the traction forces at the opposing ends of the cell, with respect to the wound direction (Eq.2). B. Snapshots of the simulated cells during an example of a recruitment process, with cells arriving in the wound (red) involved in secreting chemoattractant. C. Heatmap indicating the speed of simulated cells towards the wound at different locations, at the same times of (B). D. Time series of the accumulated number of immune cells arriving in the wound (black line) and number of bacteria (blue line) from the example in (B, C) (see Movie S3). E. Results of 40 simulations of different random initial conditions, for several values of the chemoattractant secretion strength parameter *α* (Eq.3). We plot the mean number of recruited immune cells as function of time, for the different values of *α*. F. Average radial speed towards the wound as function of time, for simulated cells outside the wound, normalized by the value of the first peak of the WT-like cells (black line, α=10). Pink line gives the data for the 5LO-like cells (α=35). G. Average radial speed towards the wound as function of time in experiments, for the cells outside the wound, normalized by the value of the first peak of the WT cells (black line). Pink line gives the data for the 5LO fish. H. Plot of the log of the number of bacteria vs the number of recruited neutrophils (at time=100min) in simulations, for 40 simulations of different random initial conditions. The scatter plot shows the results for WT-like (black) and 5LO-like (pink) cells. The crosses denote the average values with error bars for standard deviation. I. As in (H) from the experiments (recruited neutrophils at 120min, and bacteria count at 24hrs). The crosses denote the average values with error bars for standard error of the mean. Error bars represent standard error of the mean.

To simulate experimentally observed self-termination of swarming^10,11,15,23^, we implemented receptor saturation^23^ which leads to a decrease of the migration speed of immune cells towards the wound at high levels of chemokine concentrations. However, receptor saturation alone was insufficient to recapitulate the rapid arrest of swarming in previously described experimental sterile wounds (see Fig. 5C from ref^23^ compared with **Fig.S3C**). Various pathways reported in the literature could play the role of active stop signals, such as NADPH oxidase and lipoxins^11,24,54^. Therefore, we added an active stop signal, whereby cells spontaneously switch from secretion of chemokine to producing an inhibitor of chemoattractant secretion. This reproduced the rapid deceleration of swarming seen experimentally in sterile wounds in previous work^23^ (**Fig.S3C**).

**Figure 5:**
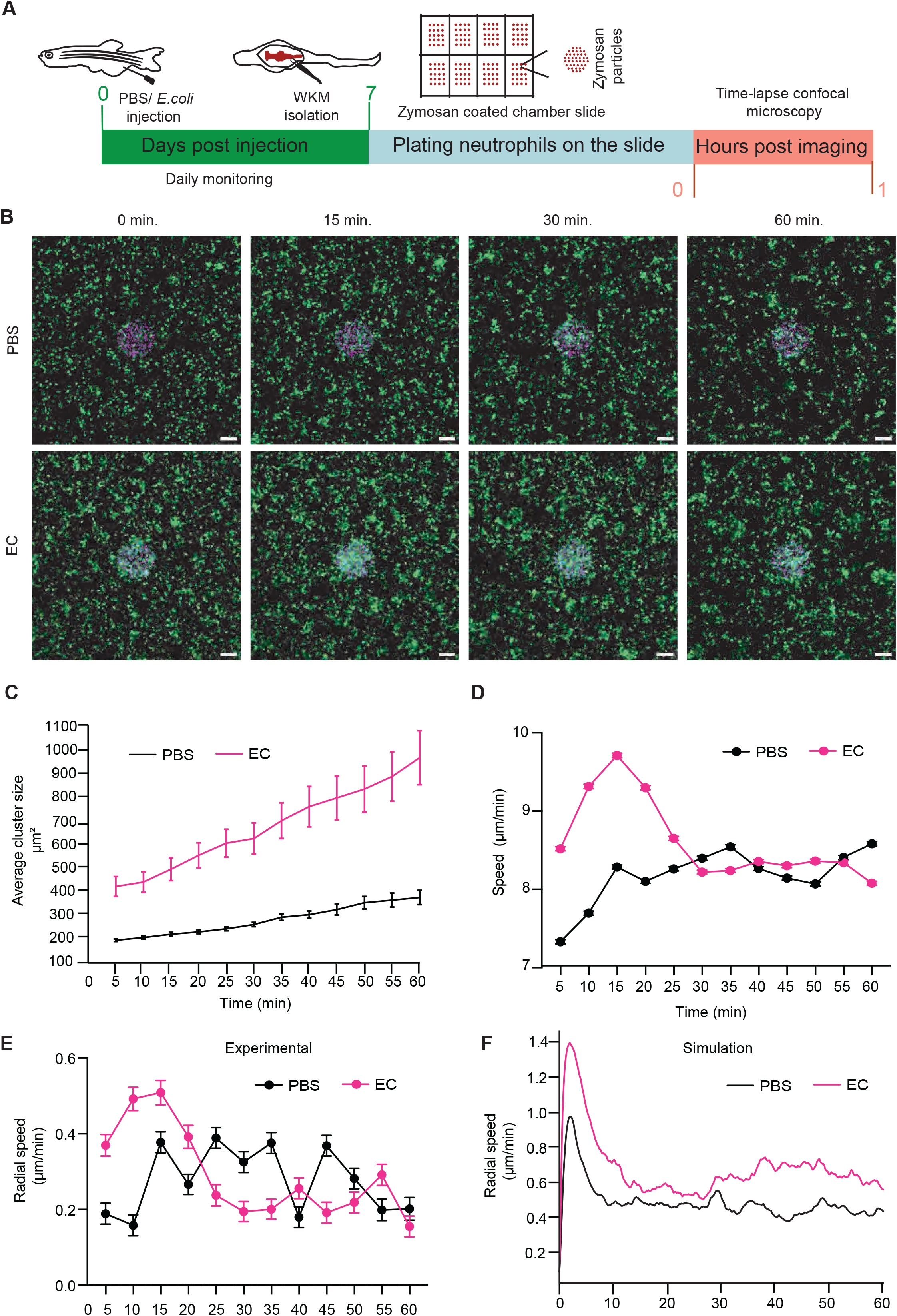
Immune training induces cell-intrinsic alterations in neutrophil swarming. A. Schematic of experiment with control, PBS pre-challenged or trained, *E. coli* pre- challenged (EC) adult zebrafish via intraperitoneal injection. Marrow collection and in vitro swarming imaging assay. B. Representative time-lapse images showing neutrophil (isolated neutrophils from adult zebrafish, seven days post PBS or *E. coli* pre-challenge) swarming towards patterned zymosan in vitro at the indicated time points. Scale bar 50 μm. C. Relative area of the neutrophil clusters overtime in control, (PBS n=10 videos from 2 independent experiments and a pool of 7 animals) (black) and EC (n=10 videos from 2 independent experiments and a pool of 7 animals) (pink) injected animals during their migration towards patterned zymosan D. Radial speed quantification of tracked cells from control (PBS n=10 videos from 2 independent experiments and a pool of 7 animals) (*black*) and EC (n=10 videos from 2 independent experiments and a pool of 7 animals) injected animals *(pink)* overtime during their migration towards patterned zymosan E. Speed quantification of tracked cells from control (PBS n=10 videos from 2 independent experiments and a pool of 7 animals) (*black*) and EC (n=10 videos from 2 independent experiments and a pool of 7 animals) injected animals *(pink)* overtime during their migration towards patterned zymosan F. Average speed towards the wound as function of time, for simulated cells outside the wound, normalized by the value of the first peak of the WT-like cells (black line, α=10) and *E.coli*-trained-like cells (pink line, *α*_0_ = 11, *α*_1_ = 89 with *B_s_* =350 (Eq.11). In this simulation the bacteria (representing the patterned zymosan) are kept at a constant value of *B* = 20 (as the zymosan does not grow as the bacteria do in the in- vivo wound and is also not eliminated by the neutrophils). Error bars represent standard error of the mean.

We then exploited this self-limiting core production model to determine the role of chemoattractant secretion rates in swarm kinetics. To explore the role of chemoattractant secretion rates, we simulated swarm dynamics with varying *α* rates (**Fig 4E-F**). This showed that the magnitude of recruitment scales with *α* (**Fig 4E**) and that the radial speed shows a higher peak during swarming **(Fig. 4F**), which fits with our experimental data from 5-LO-Neu versus control fish (**Fig 4G**). Notably, when *α* becomes too large, the strong saturation of chemotaxis diminishes the recruitment of neutrophils (**Fig.4E**). In addition, in the presence of bacteria the model predicts more persistent neutrophil recruitment (radial speed does not return to baseline-**Fig. 4F**) as compared to sterile settings (radial speed returns to baseline- **Fig. S3C**). This is because the model assumes neutrophils do not produce stop signals whilst engaging with bacteria.

Our model predicted that a higher *α* rate increases the average neutrophil number recruited to the wound and consequently reduces the bacterial burden for individual animals (**Fig. 4H**). To compare this with experimental data, we plotted the number of neutrophils recruited after 1h in individual 5-LO-Neu and control larvae, versus the average bacterial burden of the same larvae after 24h (**Fig. 4I**). Though the spread of values was much larger in experimental data, we could see that they followed a similar trend as the simulations (**Fig. 4H-I**), shifting the average of burden and scale of recruitment per larva.

The experimental data showed large heterogeneity, which could be due to biological and technical variability. However, we were intrigued to see that even in the simulations, there was considerable scatter in the values of predicted bacterial burden even within larvae with the same scale of recruitment (**Fig. S3D-F**). We hypothesised that this variation was due to differences in the initial distribution of neutrophils around the wound. To test this, we measured the average distance of neutrophils from the wound at the initial timepoint from representative examples of larvae that had different bacterial burden but similar final number of neutrophils (**Fig. S3G**). This showed that the number of immune cells at a close distance from the wound was higher in the larvae that ended up with low bacterial burden (**Fig. S3G**). This prediction matched experimental observations from representative examples of neutrophils from 5-LO Neu larvae (**Fig. S3H**).

Together, this evidence showed that a self-limiting core production model recapitulates the basic pattern of neutrophil swarming and predicts that cell intrinsic changes in secretion rates are sufficient to change the functional outcome of swarming in bacterial clearance.

#### Cell intrinsic changes in neutrophils after training improve swarming independent of nature of target

In our mathematical simulations, tuning up the secretion rate was sufficient to recapitulate the enhanced peak of neutrophil recruitment in 5-LO-Neu larvae. However, *E. coli*-trained animals also showed additional peaks of neutrophil recruitment, which were not predicted by this simple model. One possibility is that the training affects host cells other than neutrophils, which in turn affects the chemoattractant landscape. Another hypothesis is that neutrophils in these fish may undergo more complex reprogramming than with simple 5-LO engineering. Specifically, we reasonsed that the secretion of chemoattractant in trained larvae is increased in a manner dependent on the bacterial detection. To test this, we simulated neutrophils with a basal level of secretion strength *α*_0_, which can be further increased in response to bacterial presence (Eq.11 in Material and Methods). This combination of higher sensitivity to bacteria and increased secretion rates recapitulated observed kinetics of delayed enhancement of neutrophil swarming in trained fish (**Fig.S4A and B**) as well as the relationship of bacterial burden and scale of neutrophil recruitment (**Fig. S4C and D**). This suggested that the complex kinetics of neutrophil recruitment in *E. coli*-trained animals could emerge from cell-intrinsic alterations of neutrophils combined with a dynamically changing microbial target.

The prediction generated by this model was that neutrophils from trained fish would show simpler migration kinetics with a fixed and inert microbial target. To address this, we developed a new training challenge model and in vitro swarm assay with neutrophils from adult zebrafish. We challenged adult zebrafish with either PBS or *E. coli*, then isolated neutrophils seven days later and seeded them on slides with patterned zymosan, shown previously to potently stimulate swarming of human neutrophils^15^ (**Fig. 5A**). We found that neutrophils from *E.coli-*trained animals showed larger clustering at the target (**Fig. 5B and C, Movie S4**) and faster chemotaxis (**Fig. 5D and E, Movie S4**) compared with neutrophils from PBS-challenged animals. To confirm the validity of our mathematical model with these conditions, we employed simulations using a fixed bacterial population (where bacteria numbers cannot grow or be eliminated). We found that under a fixed target, our simulations predicted the enhanced peak of recruitment observed in vitro **(Fig 5. F)**, and no delayed peak as observed in vivo, thereby supporting our model for neutrophils from *E. coli-*trained animals.

Together, this evidence indicates that cell instrinsic alterations of neutrophils in trained animals are sufficient to explain the altered neutrophil kinetics in vivo.

## Discussion

Here, we examine whether self-generated guidance in neutrophil migration is a programmable process and the extent to which such reprogramming can affect disease outcome. We show that neutrophil swarming and the expression of genes for swarm machinery are altered during natural reprogramming (‘training’) by microbial experience. Through chemical inhibition, we found that these alterations contribute to enhanced resistance of trained animals to pathogenic infection. Through quantitative live imaging and modelling, we demonstrate how variations in secretion of chemoattractant by neutrophils at the wound lead to changes in swarm kinetics, bacterial clearance and infection outcome. Finally, we show that enhancing the ability of neutrophils to produce autologous chemoattractants (through 5-LO engineering) is sufficient to accelerate neutrophil swarming and improve bacterial clearance and disease progression.

Our study highlights novel routes for training in wound infection settings. We observed improved resistance to virulent bacterial wound infection after a pre-challenge with non-pathogenic *E. coli*. Probiotic *E. coli* strains have been reported to have beneficial effects on viral resistance in mouse models^55^, but such training has not been reported for wound infections. Neutrophil training in zebrafish larvae arises through changes in hematopoietic stem cells within similar timeframes post-infection, as has been reported in adult mammals^40^. Our observations are consistent with central reprogramming of neutrophils, as the protective effects were seen across heterologous tissue sites and the gene expression changes were detectable in neutrophils isolated from the hematopoietic tissue. In addition, we showed that zebrafish neutrophils undergo epigenetic modifications on swarm-associated genes during training with *Shigella flexneri* (which are phylogenetically close to *E. coli)* in a comparable timeframe with our study^42^.

We found that neutrophil swarm capacity is amenable to reprogramming after microbial experience. Specifically, neutrophils upregulated expression of genes associated with swarming and trained larvae were more likely to show potent swarming and low susceptibility to infection. Interestingly, we found that *E. coli* training also reduced the higher-end values for neutrophil numbers at the wound. One potential explanation is that the effect of increasing neutrophil numbers has non-linear effects on bacterial clearance, whereby at low neutrophil numbers the changes in neutrophil kinetics can make a significant difference in bacterial clearance whereas beyond a certain number of recruited neutrophils, the benefit decreases. Our mathematical modelling supports this model and suggests that beyond a certain point, increasing the secretion of chemoattractant has no benefit for bacterial clearance and can even be detrimental to microbial eradication. A question raised is why neutrophil swarming has not evolved to be maximal at steady state. In our model, enhanced swarming did not delay regeneration, suggesting that there is no detriment to physiological wound repair processes. We therefore speculate that adaptive reprogramming of neutrophil swarming may reflect a strategy to economise energy or a means to balance the risks of overt inflammation in chronic inflammatory settings.

We used mathematical modelling to understand how the dynamics of chemoattractant secretion influence neutrophil swarm kinetics and bacterial clearance. Our simplified model shows that core chemoattractant production is sufficient to generate a primary level of clustering and capture observed migration kinetics. This agrees with in vivo mouse data whereby optogenetic activation of calcium in a core cluster of cells was sufficient to generate moderate clustering^22^. It remains a future challenge to expand this model to larger-scale swarm organisation and implement additional mechanisms, such as chemoattractant relays. More knowledge of the dynamics of chemoattractant secretion will assist in that direction, for example through further development of biosensors^56^. A complexity to account for is that the same attractant (LTB4) appears to be involved in both core secretion^10^ and active relays^11^. Therefore a key question to resolve for future models is what mechanistic rules determine whether swarms limit themselves to core secretion or escalate further through signal relays.

Our experimental data indicated distinct kinetics of swarm enhancement in trained animals versus 5-LO engineered animals, with a delayed peak of swarming in the former and an early spike in the latter. A general explanation is that in trained animals a range of genes are altered whereas in 5-LO engineered animals, the enhancement is specific to chemoattractant secretion. We explored theoretical parameters that could be altered during training. Our mathematical model predicts that a combination of upregulation of swarm machinery (e.g. 5-LO expression) together with enhanced sensitivity to bacterial detection and bacteria-dependent secretion can produce the delayed swarm peak as seen in trained fish. This theory aligns with evidence from the literature, in that LTB4 secretion is triggered by bacterial detection through Toll like receptor (TLR) signalling^57^ and that TLR signalling pathways can be reprogrammed during trained immunity^32^. It is noteworthy that epigenetic H3K4 marks were detected along the catabolic regulator of LTB4 *cyp4f3*, suggesting that training may fine tune LTB4 dynamics in a more complex manner. Future studies could aim to dissect these circuits and recapitulate these by genetic engineering.

Our discovery that engineering 5-LO is sufficient to improve bacterial clearance has relevance to cellular immunotherapies. Strategies for genetic modification of immune cells are growing and extending to cell types beyond lymphocytes, due to advances in genetic engineering and stem cell biology^44,45,47,58^. In this backdrop, the engineering of chemoattractant receptors has been conceptualised to improve the efficacy of CAR-engineered immune cell deployment to tumors^59^. Our study suggests that harnessing the ability of immune cells to seed chemoattractant gradients in target tissues may offer new routes for reprogramming immune cell migratory patterns.

## Acknowledgements

The *P. aeruginosa* strain was kindly provided by Prof. Martin Welch, Department of Biochemistry, University of Cambridge and the *E. coli* strain was kindly provided by Dr. Somenath Bakshi, Department of Engineering, University of Cambridge. We thank Christian Godfrey and Seok Joon Mun at the BioMEMS Core at the Massachusetts General Hospital for providing the swarm-array slides. We thank Margarida C. Gomes and Serge Mostowy for sharing the ChIP-seq data from trained zebrafish. S.M.B., I.W., G.R., C.C., Y.L. were supported by the European Research Council (ERC) under the Horizon 2020 program and UKRI, Grant agreement No. EP/Y02799X/1. Y.L. is also supported by a BBSRC DTP programme BB/X010899/1. N.S.G. is the incumbent of the Lee and William Abramowitz Professorial Chair of Biophysics (Weizmann Institute) and acknowledges support from the Royal Society Wolfson Visiting Fellowship (RSWVF\R2\2320010), Human Frontier Science Program grant RGP0032/2022 and ISF-DFG grant 1182/25. H.M.C. was supported by the Roving Researcher scheme in the School of Biological Sciences, University of Cambridge. The authors gratefully acknowledge the help by the Aquatics facility at the Department of Physiology, Development and Neuroscience, and the Microscopy Bioscience Platform, University of Cambridge for their support and assistance in this work.

## Competing Interests

The authors declare no competing interests

## Code availability

The available codes used for modelling neutrophil swarming can be found here: https://github.com/samantatuhin1991/immune-swarming-model

## Methods

### Experimental model and subject details

All zebrafish were maintained in accordance with UK Home Office regulations under the UK Animals (Scientific Procedures) Act 1986 (PPL number PP4458156), with approval from the University Biomedical Service Committee. Adult fish were maintained and bred following established protocols^60^. Briefly, zebrafish were bred and maintained under standard conditions at 28.5 ± 0.5 °C on a 14 h light:10 h dark cycle. Embryos were collected from natural spawnings at 3 hours post-fertilization (hpf), bleached for 5 minutes in 0.003% sodium hypochlorite (NaOCl; Cleanline, CL3013), and rinsed three times with E3 medium. They were subsequently incubated at 28 °C in E3 medium supplemented with 0.1 µg /ml methylene blue (Sigma-Aldrich, M9140-25G). For imaging experiments, E3 medium was additionally supplemented with 0.003% 1-phenyl-2-thiourea (PTU; Sigma Aldrich, P7629) to inhibit pigmentation and preserve optical transparency.

For experiments that required *E. coli* training, *E. coli* was administered to larvae at 2 days post-fertilization (dpf) and subsequent re-challenge experiments were done at the 4 dpf stage. In experiments without training, larvae at 3-4 dpf were used. For live imaging, larvae were anesthetized in appropriate buffer containing 160-200 mg/L MS-222 (Sigma-Aldrich, E10521- 50G). Where indicated, larvae were treated with Zileuton (20 μM, Tocris, 3308) or appropriate volume of vehicle (DMSO (Sigma-Aldrich, 276855-100ML)) solution by immersion.

The following transgenic zebrafish lines were used: Tg(*lyz*:GCamp6F)*^cu^*^104^ and Tg(*lyz:*tRFP- 5LO) *^cu1^*^06^ from previously described work^10^,Tg(*mpx*:GFP)i114^61^, Tg(*runx1+23*:mCherry).

### Bacterial culture and preparation

For *E. coli* injection, cyan fluorescent protein (CFP)-expressing *E. coli* MG1655 7740 strain was developed as described previously^62^. A single colony was obtained from a four-way streak on antibiotic-free LB Agar (Melford, A20020-500.0) plate. The colony was inoculated into 50 ml of antibiotic-free LB broth (Formedium, LBX0102) and cultured at 37 °C with shaking until the OD_600_ reading reached 0.8-0.9. The culture was then aliquoted into 1 ml stocks in antibiotic-free LB broth with 20% v/v glycerol (Fisher Scientific, G/0650/17), snap-frozen with liquid nitrogen, and stored at -80 °C until usage. On the day of the experiment, aliquots of *E. coli* were thawed at 37 °C, pelleted by centrifugation at 5,000 × *g* for seven minutes at 4 °C, and washed twice with PBS (Sigma Aldrich, D8537). The bacteria were finally suspended in PBS with 10% v/v phenol red (Sigma, P0290).

For *P. aeruginosa* infection, PAO1 strain was used^10^. One day prior to infection, a single colony of *P. aeruginosa* was obtained from a four-way streak on Pseudomonas Isolation Agar (PIA; BD Difco, 292710) supplemented with cetrimide and nalidixic acid (E&O Laboratories Ltd, LS0006) and glycerol. The colony was inoculated into 5 ml of antibiotic-free LB broth and cultured at 37 °C with shaking for 24 h. On the day of the experiment, the overnight culture was diluted in fresh LB and incubated to mid-logarithmic phase (OD₆₀₀ = 0.6–0.8). Bacterial density was adjusted to 3 × 10⁵ CFU/mL using spectrophotometric estimation. Bacteria were pelleted at 1700 × *g* for five minutes and washed twice with PBS to remove residual growth medium, then resuspended in PBS.

### PBS and *E. coli* injections

On the day of injections, *E. coli* injection mix was prepared as described above. PBS injection mix was prepared as PBS with 10% v/v phenol red for control. Larvae were anaesthetised in E3 medium supplemented with 0.1 µg/ml methylene blue and 160-200mg/L MS-222. The left otic vesicle of each zebrafish larva was then injected with 2 drops of 0.5 nL of PBS or *E. coli* injection mix with a pressure of 20 psi using Picospritzer III microinjector (Intracell), as previously described^63^. To confirm the dose of *E. coli* injected, 2 drops of 0.5 nL of injection mix was directly injected into a microcentrifuge tube containing 100 µl PBS after the injections and were then plated onto antibiotic-free LB Agar plates overnight at 37 °C at 10^0^ and 10^-1^ dilution. The bacterial colony forming units (CFU) were then determined by counting the number of CFP^+^ colonies after overnight incubation. Only when the *E. coli* injection dose exceeded 400 CFU/ml were the larvae included for subsequent experiments. After injections, larvae were transferred to fresh E3 medium supplemented with 0.1 µg/mL methylene blue and kept at 28 °C until further use.

For adult training experiments, zebrafish Tg(mpx:GFP)i114 were anaesthetised with 200 mg/L of MS-222 until unconscious. 5 uL of either PBS or a solution of 4 x 10^6^ CFU in PBS of *E. coli* were injected in the peritoneal cavity. The fish were immediately placed in the recovery tank and monitored until full recovery. The injected fish were kept for 7 days post injection and monitored twice daily.

### Tail transection, wound infection and survival assay

Zebrafish larvae were transferred to a Petri dish containing Ringer’s solution and 160-200 mg/L MS-222. Tail transection was done using a surgical blade (Swann Morton, 0510) at the distal boundary of the notochord, avoiding injury to the notochord. Non-transected larvae served as controls. Immediately after tail transection, larvae were transferred to a six-well plate containing Ringer’s solution with 3 × 10⁵ CFU/mL *P. aeruginosa* and incubated at 32 °C for 3 h. After incubation, larvae were washed five times in E3 medium (without methylene blue). Individual larvae were then placed in wells of a 24-well plate containing 1 mL E3 (without methylene blue) and monitored for survival and disease progression. For 5-LO inhibition assays, 20 μM zileuton treatment (diluted in DMSO) was initiated 1 h prior to tail transection and maintained until the end of the experiment.

Survival was assessed immediately after washing, at 3 hpi, and 24 hpi. Assessment was also done at intermediate time points (7 hpi and 10 hpi) when experiment was done for 24 h. For 48h time point experiments, analysis was done at 3 h, 24 h and 48 h. Phenotypic disease scoring was based on behavioural responses, transparency, tissue integrity, circulation, and survival. Between observation timepoints, larvae were returned to 32 °C. A score of 4 was defined as death of the organism that was determined by the absence of heartbeat as observed under the microscope.

### Measurement of regeneration after sterile tail transection

Caudal fin transection of zebrafish larvae (3dpf) was performed using a sterile surgical blade at the boundary of the notochord, without injuring the notochord. Following injury, larvae were rinsed with E3 medium and placed in individual wells of a 24-well plate in E3 medium, at 28.5°C. Images of each larva was acquired on a stereomicroscope (Zeiss Stemi 2000-C) immediately after caudal fin transection (0h), one day after transection (24h) and two days after transection (48 h).

For every larva, the caudal fin area posterior to the notochord was outlined using the Freehand selection tool in ImageJ/Fiji for images acquired at all the three time points. The outlined area was measured to obtain a total surface area of the region of interest (ROI). The area of the ROI at 24 h and 48 h for each larva was normalized by the area of the ROI for the same larva at 0 h.

### Bacterial burden quantification

At indicated timepoints, zebrafish larvae from each condition were pooled into groups of three larvae and homogenised in 300 μL PBS containing 0.1% v/v Triton X-100 (VWR, 28817.295) using sterile pestles. Serial dilutions (up to 10⁻³) were prepared in PBS and plated on PIA supplemented with cetrimide and nalidixic acid (as per manufacturer’s instructions) for *P. aeruginosa* or antibiotic-free LB agar for *E. coli.* CFU were then counted after 24 hours incubation.

### Fluorescence-Activated Cell Sorting (FACS)

Approximately 400 embryos from a cross between Tg(*mpx*:GFP)*^i1^*^14^ and Tg(*runx1+23*:mCherry) transgenic lines were injected with either *E. coli* or PBS (200 per group) in the otic vesicles at 2 dpf as described above. At 4 dpf, larvae were collected into microcentrifuge tubes, rinsed twice with 1x HBSS (prepared by diluting 10x HBSS (Gibco™, 14175095) in UltraPure™ DNase/RNase-Free Distilled Water (Invitrogen™, 10977035)), and placed on ice for 10 minutes to immobilise the larvae. The caudal haematopoietic tissue (CHT) regions from the larvae were dissected on sterile petri dishes using ethanol-sterilised scalpels and centrifuged to remove excess HBSS.

For tissue dissociation, 500 μL of digestion solution (2% w/v collagenase (Gibco™, 17104019) and 40% v/v trypsin (Gibco™, 15400054) in 1x HBSS) was added to each tube. Samples were incubated at 30 °C for 30 minutes with periodic trituration by pipetting. Digestion was halted with 1x HBSS containing 10% v/v heat-inactivated fetal bovine serum (FBS; Gibco™, A5670701). All subsequent steps were performed on ice. Cells were centrifuged at 500 × g for 5 minutes at 4 °C and then resuspended in sorting solution (1x HBSS supplemented with 10 mM HEPES 15630056) and 2% FBS) and filtered through a pre-wetted 40 µm cell strainer (Falcon, 352340). Filtered cells were centrifuged and resuspended in 1 ml sorting solution in a microcentrifuge tube. Cell viability and concentration were assessed using trypan blue (Gibco, 15250) staining and a hemocytometer, respectively.

GFP-positive neutrophils were FAC-sorted using a BD FACS Aria III (Becton Dickinson) directly into 100 μL of lysis solution supplied with the RNAqueous® Micro Kit (ThermoFisher, AM1931).

### RNA extraction and cDNA synthesis

For transcriptomic analysis, RNA isolation was done from the sorted cells using the RNAqueous® Micro Kit (Thermo Fisher, AM1931) according to the manufacturer’s instructions, including DNase I digestion step to eliminate genomic DNA contamination.

For cytokine expression analysis following PBS/ *E. coli* injections, at 2 dpf, RNA was extracted from groups of 10 wild-type zebrafish per condition (non-injected, PBS-injected, and *E. coli*- injected) after euthanasia with MS-222 at 4.5 hours post-injection using the RNeasy® Mini Kit (Qiagen, 74104). Subsequently, from 3 to 10 dpf, groups of 8-10 larvae per condition were euthanised with MS-222 prior to RNA extraction.

The obtained RNA concentrations were measured using the Nanodrop. Purified RNA was then reverse transcribed into cDNA using the Maxima H Minus First Strand cDNA Synthesis Kit (ThermoFisher, K1651) following the manufacturer’s protocol. Finally, cDNA was stored at - 20 °C until further use for quantitative PCR (qPCR) analysis.

### Gene expression analysis

The qRT-PCR was performed using the PowerTrackTM SYBR^TM^ Green Master Mix (ThermoFisher, A46012) and carried out in the LightCycler® 480 Instrument II (Roche). For transcriptomic analysis, each reaction contained 0.5 ng of cDNA, 10 nM of primer pairs, and 1X master mix. The zebrafish gene *rpl32* served as the reference for normalisation.

For cytokine expression analysis, each reaction contained 5 ng cDNA, 500 nM primer pairs, and 1X master mix and the zebrafish gene *ef1a* served as the reference for normalisation. The mRNA levels of target genes relative to a housekeeping gene (**Table 1)** were calculated using the Livak (2-^−ΔΔCqt^) method^64^. All qRT-PCR reactions for transcriptomic analysis were performed in duplicate with five biological repeats for transcriptomic analysis while the cytokine expression experiments were performed in triplicate with three biological repeats.

**Table 1.**
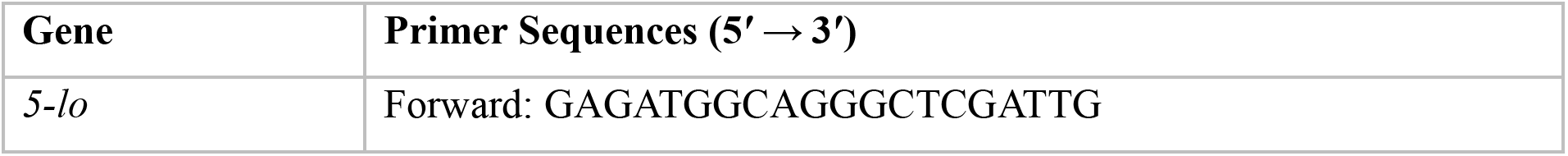
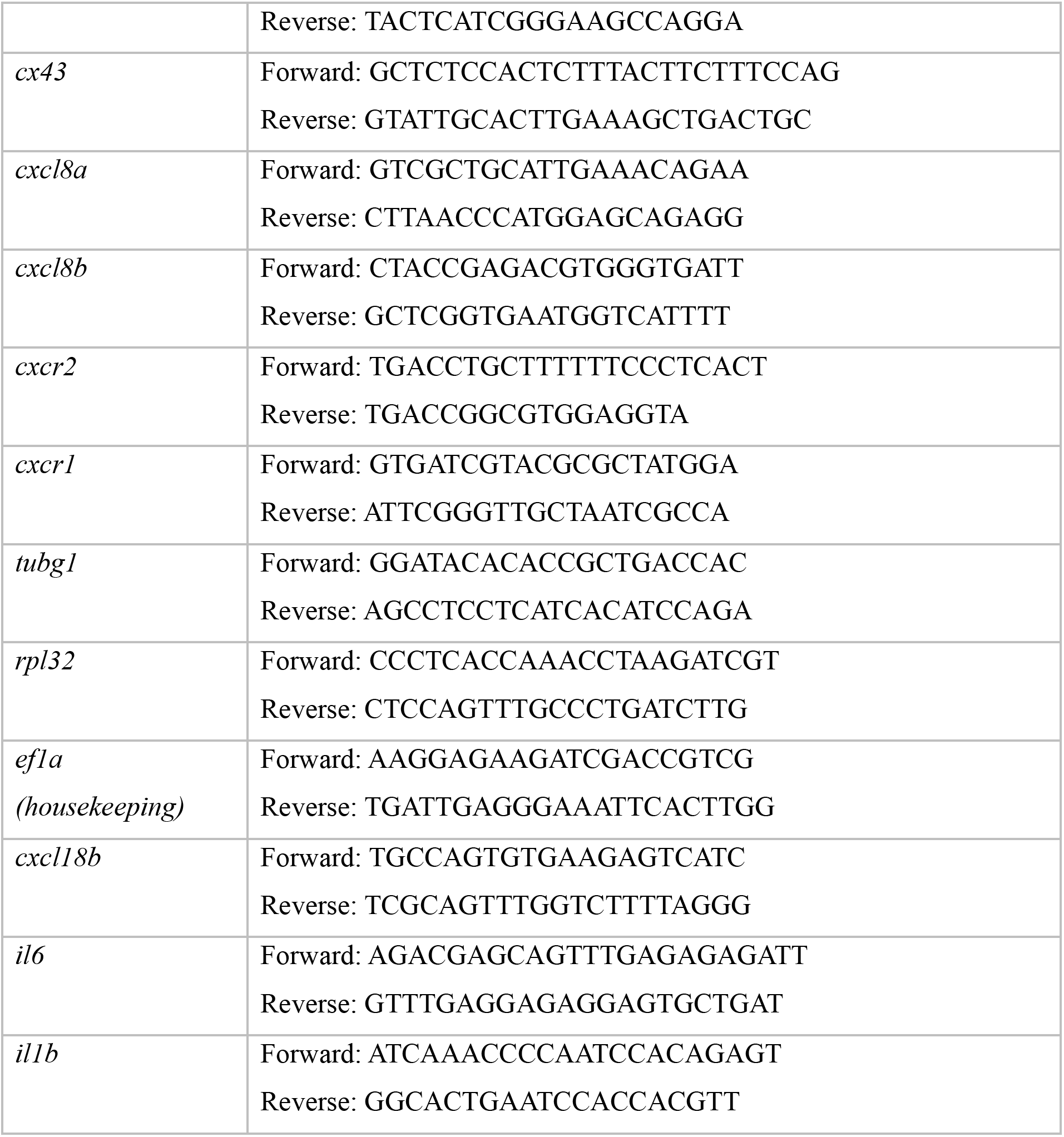
ChIP-seq Data Analysis and Visualisation. H3K4me3 ChIP-seq data from *Shigella*-trained zebrafish neutrophils were obtained from GEO accession GSE217063^42^.Processed peak files were filtered to extract signal values and q-values for promoter regions (≤1 kb from transcription start site) of five swarming-associated genes (*cyp4f3, lta4h, alox5a, pla2g4aa, alox5ap*). Heatmaps were generated using the pheatmap package in R, with statistical significance (q < 0.05) indicated by asterisks. Permission for data use was obtained from the corresponding author of the original study.

### Two-photon imaging and laser wounding

Two photon imaging was performed in Tg(*lyz*:GCaMP6f)*^cu104^*larvae or larvae from crosses between Tg(*lyz*:GCaMP6f) x Tg(*lyz:*tRFP-5LO). Transgenic larvae expressing the calcium indicator GCaMP6f were screened to ensure high reporter expression. Larvae exhibiting strong fluorescence and normal morphology were selected for imaging and infection. Selected larvae were anaesthetised and mounted in a mixture of 1.2% low-melting-point agarose (Invitrogen, 16520) and the medium (Ringer) containing 160-200 mg/L MS-222. Imaging under sterile conditions was performed as as previously described^65^. For sterile imaging, larvae were mounted on a 35 mm glass-bottom dish with a 14 mm microwell #1.5 coverglass. Excess agarose was removed before solidification to embed the larvae in a thin layer. For infection imaging, larvae were mounted in a custom-built coverslip chamber consisting of glass coverslips sealing both the top and bottom of the sample onto a metallic ring. Larvae were oriented laterally, and the agarose was allowed to solidify. After agarose solidification, the urogenital region was exposed by carefully removing surrounding agarose under a dissecting microscope using a pair of forceps, allowing bacterial access post-wounding. The chamber (for infection conditions) or dishes (for sterile conditions) were filled with 1 mL or 4 mL of Ringer’s solution containing 160-200 mg/L MS-222 and 3 × 10⁵ CFU/mL PAO1. For sterile conditions, appropriate Ringer buffer was used as indicated.

Laser ablation and time-lapse imaging were performed on a LaVision TriM Scope multiphoton microscope equipped with an electro-optic modulator for rapid power modulation. A Spectra-Physics Insight DeepSee dual-line laser was tuned to 900 nm for imaging and 1,040 nm for ablation, with imaging power set to ∼500 mW at the specimen plane. Image acquisition was controlled using ImSpector Pro software (version 5.0.284.0; LaVision Biotec, ©1998–2016). Two-photon imaging was conducted with a 25×/NA 1.05 water-dipping objective. A 40 μm- diameter region of interest was defined on a single superficial focal plane (240 nm/pixel, 15 μs dwell time). Z-stacks (∼20 planes, 2 μm step size) were acquired every 20 s. Laser wounding was initiated after a 2 min pre-wound baseline, followed by 2 h of post-wound imaging. Focus was maintained throughout acquisition. Image stacks were processed in Fiji using maximum-intensity z-projections to generate neutrophil-swarming time-lapse movies for subsequent analysis.

#### Adult marrow collection and *in vitro* swarming assay

Following training, on the 7th day after injection Schedule 1 procedure, euthanasia with overdose of MS-222 and destruction of the brain, was performed.

Marrow tissue was collected under a fluorescent microscope by incision on the fish and removing all the internal organs. Marrows were collected in a Eppendorf with 1 mL of HBSS on ice^66^. The collected tissue was then filtered through a 40 uL strainer to create a single cell solution. The cells were resuspended in a total of 6 mL of HBSS and gently layered on top of 2 mL of density gradient leukocyte separation medium 1078 (Mediatech; CellGro, AK) in a 15 mL falcon tube and centrifuged (400 g, 30 min) (Isles et al., 2021). The layer of leukocytes was collected in a 15 mL falcon tube in a total volume of 4 mL of HBSS and centrifuged (400 g, 15 min). The total number of neutrophils was defined by hemocytometer counting, under a fluorescent microscope. Neutrophils were resuspended at 3 x 10^6^/mL in RPMI + 10% FBS and 200 uL were placed in the zymosan swarming chamber^67^. Imaging was started as soon as possible. Time lapses were acquired at 2 location per condition. Images were analysed as followed in the next section. The neutrophil swarming chambers were manufactured at the BioMEMS Core at the Massachusetts General Hospital in Boston https://researchcores.partners.org/biomem/about .

#### Image analyses

##### Defining the cumulative number of cells arriving at the wound

The cumulative number of neutrophils that reach the wound were calculated by manually counting the neutrophils reaching the wound in blinded datasets using ImageJ/Fiji overtime.

##### Cell trajectories speed and radial speed

For *in vivo* experiments, neutrophils were semi-automatically tracked using Imaris v8.2 (Bitplane AG, Zürich, Switzerland) on 2D maximum intensity projections of the 4D time-lapse movies. For *in vitro* experiments videos were uploaded into imaris software, a square of 410 um^2^ was drawn and the GFP channel was used to track the cells overtime. Cell position data of the tracked cells overtime from Imaris were exported using Excel 2016. Speed and radial speed were calculated using MATLAB scripts developed previously^10^. Briefly, for radial speed calculations, instantaneous speed values overtime for individual neutrophils were multiplied for the cosine of the angle. 5 minutes bin average was performed. In *in vivo* experiments tissue drift was observed in the first 20 minutes after LW, therefore, these timepoints were excluded from speed and radial speed analyses. Movies in which there was considerable tissue drift throughout the imaging window or where the wound was too small to cause any detectable response were excluded from the analysis (first 25 min).

##### Cluster area measurements

A region of interest of 200 *μ*m^2^ was drawn per video around the zymosan target. Imaris’ surface detection feature of was used to measure the cluster area overtime, and the data was exported in Excel. The average cluster area per timepoint per video and condition was calculated and plotted.

## Statistical analyses

All statistical tests were performed in Prism10 (GraphPad Software, La Jolla, CA). The statistical test and the n number are indicated in the figure legends. The error bars show standard error of the mean except for bacterial burden data where error bars represent 95% confidence intervals of the median. Where the distribution was verified as normal, outliers were removed by applying Rout test. Live imaging experiments were acquired in minimum five independent experiments. A normal distribution test was performed prior to any *t*-test statistical analyses. Normal distribution was assessed using *Shapiro-Wilk test*.

### Theoretical model

The signal triggered by the local concentration of the chemokine (c(r)), at the cell edge is given by (Fig.5(a,b)):

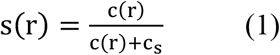

And the net cell speed up the chemokine gradient is given by the difference of the traction forces across the opposite ends of the cell, with respect to the direction of the wound (given by unit vector *n*\*, Fig.5(a,b))

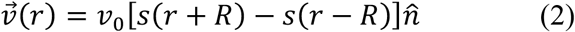

Where *R* is the typical cell radius and *v*_0_ its maximal migration speed. We ignore here the random component to the cell speed, so that we focus on the average speed towards the wound due to the chemokine signal.

We focus here on the long-time swarming behavior, which we assume to be dominated by the chemokine signal that is secreted at the wound. We assume that the chemokine concentration has an exponentially profile decaying radially away from the wound, with its amplitude at the wound cite dependent on the number of chemokine secreting neutrophils (*n_s_*). Neutrophils arriving at the wound are assumed to start secreting chemokine, so that the chemokine profile has the form:

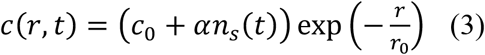

The equation for the evolution of the number of chemokine secreting neutrophils at the wound is given by

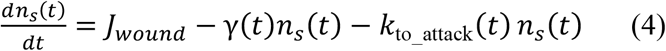

Where *J*_w*ound*_ is the flux of cells arriving at the wound from the migration domain (Fig.Xa). The second and third terms describe the rates at which the chemokine secreting cells change their states to “stop-signal” mode (*n_ss_*) and bacteria-attack model (*n_b_*).

The secreting neutrophils undergo a spontaneous transition to a “stop-signal” mode (*n_ss_*), at a rate *γ*. We assume that this rate has a basal value (*γ*_0_), but is dominated by the “stop-signal” secreted by *n_ss_* cells, in the form:

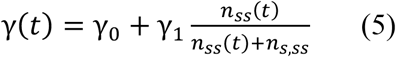

Where *γ*_1_ is the parameter describing the “stop-signal” effectiveness, and *n_s_*_,*ss*_ describes the saturation of the response to this signal. The number of “stop-signal” cells evolves according to the following equation

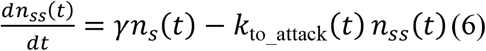

Note that while the influx of cells into the wound is discrete, determined by the arrival of the particles migrating to the wound within the simulation domain (Fig.5a), the subsequent evolution of the numbers of cells in each of the states evolve according to continuum equations (Eqs.4,6) for simplicity.

Finally, we consider the dynamics of the invading bacteria and their effect on the neutrophils. The neutrophils are able to sense the presence of bacteria and this signal, which is proportional to the number of bacteria, triggers the transition of neutrophils from the chemokine and stop-signal secreting states into “attack” mode (*n_b_*), at a rate given by

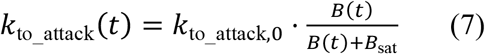

Where *k*_to_attack,0_ is some constant rate, *B*(*t*) is the number of bacteria in the wound at time *t* and *B*_sat_ is some saturation constant. The equation of the time evolution of the number of neutrophils in “attack” mode is given by

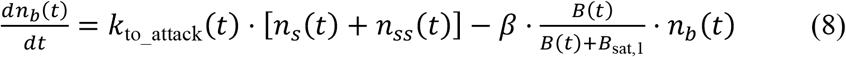

Where the last term describes the rate at which neutrophils get killed/consumed due to their engagement with the bacteria. This term includes a saturation which describes a maximal number of bacteria that each neutrophil can interact with, of order of the saturation constant *^B^*sat,1^.^

The dynamics of the bacteria is given by the following equation

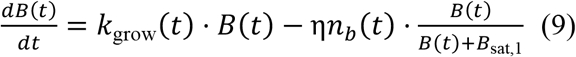

Where the first term describes their growth rate, given by

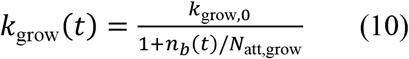

Where *k*_grow,0_ is a constant, and we consider that the neutrophils can diminish the bacterial growth rate, as described by the denominator. The last term in Eq.(9) describes the rate at which bacteria are removed/destroyed by the neutrophils.

Finally, we also consider that the secretion of the chemoattractant can be increased in response to the presence of bacteria. We use a form as in Eq.(7), such that

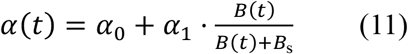

The list of the model parameters is given in Table I in the Supplmentary Information section.

This preprint originally included additional data, which have now been moved to a separate preprint. **doi:** https://doi.org/10.64898/2026.08.04.742764

## Notes

### Competing Interest Statement

The authors have declared no competing interest.

### Summary of Updates

This revision includes new data in Figure 5. In addition, we have removed data pertaining to Figure 1 of the original version and included these datasets in a new preprint (https://doi.org/10.64898/2026.08.04.742764). The authorship list has been updated to reflect the split contributions. We have added a footnote in both preprints on the first page of the manuscript to explain the relationship of the two preprints.

## References

1. Schrope, J. H., Robertson, T. F., Sarris, M. & Huttenlocher, A. Chemical and Mechanical Regulation of Leukocyte Migration. Cold Spring Harb. Perspect. Biol. a041752 (2025) doi:10.1101/cshperspect.a041752.

2. Insall, R. H., Paschke, P. & Tweedy, L. Steering yourself by the bootstraps: how cells create their own gradients for chemotaxis. Trends Cell Biol. 32, 585–596 (2022).

3. Stock, J., Kazmar, T., Schlumm, F., Hannezo, E. & Pauli, A. A self-generated Toddler gradient guides mesodermal cell migration. Sci. Adv. 8, eadd2488 (2022).

4. Uçar, M. C., Alsberga, Z., Alanko, J., Sixt, M. & Hannezo, E. Self-generated chemotaxis of mixed cell populations. Proc. Natl. Acad. Sci. 122, e2504064122 (2025).

5. Dalle Nogare, D. & Chitnis, A. B. A framework for understanding morphogenesis and migration of the zebrafish posterior Lateral Line primordium. Mech. Dev. 148, 69–78 (2017).

6. Muinonen-Martin, A. J. et al. Melanoma Cells Break Down LPA to Establish Local Gradients That Drive Chemotactic Dispersal. PLoS Biol. 12, e1001966 (2014).

7. Tweedy, L. et al. Seeing around corners: Cells solve mazes and respond at a distance using attractant breakdown. Science **369**, eaay9792 (2020).

8. 8. Alanko, J., et al. CCR7 acts as both a sensor and a sink for CCL19 to coordinate collective leukocyte migration. Sci. Immunol. 8, eadc9584 (2023).

9. Lämmermann, T. et al. Neutrophil swarms require LTB4 and integrins at sites of cell death in vivo. Nature 498, 371–375 (2013).

10. Poplimont, H. et al. Neutrophil Swarming in Damaged Tissue Is Orchestrated by Connexins and Cooperative Calcium Alarm Signals. Curr. Biol. 30, 2761–2776.e7 (2020).

11. Strickland, E. et al. Self-extinguishing relay waves enable homeostatic control of human neutrophil swarming. Dev. Cell 59, 2659–2671.e4 (2024).

12. Afonso, P. V. et al. LTB4 Is a Signal-Relay Molecule during Neutrophil Chemotaxis. Dev. Cell 22, 1079–1091 (2012).

13. Dowdell, A., Paschke, P. I., Thomason, P. A., Tweedy, L. & Insall, R. H. Competition between chemoattractants causes unexpected complexity and can explain negative chemotaxis. Curr. Biol. 33, 1704–1715.e3 (2023).

14. Burn, G. L., Foti, A., Marsman, G., Patel, D. F. & Zychlinsky, A. The Neutrophil. Immunity 54, 1377–1391 (2021).

15. Reátegui, E. et al. Microscale arrays for the profiling of start and stop signals coordinating human-neutrophil swarming. *Nat*. Biomed. Eng. 1, 0094 (2017).

16. Ng, L. G. et al. Visualizing the Neutrophil Response to Sterile Tissue Injury in Mouse Dermis Reveals a Three-Phase Cascade of Events. J. Invest. Dermatol. 131, 2058– 2068 (2011).

17. Chtanova, T. et al. Dynamics of Neutrophil Migration in Lymph Nodes during Infection. Immunity **29**, 487–496 (2008).

18. Hopke, A. et al. Neutrophil swarming delays the growth of clusters of pathogenic fungi. Nat. Commun. 11, 2031 (2020).

19. Saraswat, D. et al. Neutrophil swarming is crucial for limiting oral mucosal infection by *Candida albicans*. J. Leukoc. Biol. 117, qiae239 (2025).

20. Herrero-Cervera, A., Soehnlein, O. & Kenne, E. Neutrophils in chronic inflammatory diseases. Cell. Mol. Immunol. 19, 177–191 (2022).

21. Isles, H. M. et al. Pioneer neutrophils release chromatin within in vivo swarms. eLife 10, e68755 (2021).

22. 22. Khazen, R., et al. Spatiotemporal dynamics of calcium signals during neutrophil cluster formation. Proc. Natl. Acad. Sci. 119, e2203855119 (2022).

23. Kienle, K. et al. Neutrophils self-limit swarming to contain bacterial growth in vivo. Science 372, eabe7729 (2021).

24. Song, Z., Bhattacharya, S., Clemens, R. A. & Dinauer, M. C. Molecular regulation of neutrophil swarming in health and disease: Lessons from the phagocyte oxidase. iScience 26, 108034 (2023).

25. Sun, D. & Shi, M. Neutrophil swarming toward Cryptococcus neoformans is mediated by complement and leukotriene B4. Biochem. Biophys. Res. Commun. 477, 945–951 (2016).

26. Arya, S. B. et al. cPLA2 α targeting to exosomes connects nuclear deformation to LTB4 -signaling during neutrophil chemotaxis. Sci. Adv. 12, eaea2784 (2026).

27. Strickland, E. D. et al. Arachidonic acid availability controls neutrophil swarm initiation and scaling. Preprint at 10.64898/2026.04.15.718800 (2026).

28. Hopke, A. et al. Transcellular biosynthesis of leukotriene B4 orchestrates neutrophil swarming to fungi. iScience 25, 105226 (2022).

29. Uderhardt, S., Martins, A. J., Tsang, J. S., Lämmermann, T. & Germain, R. N. Resident Macrophages Cloak Tissue Microlesions to Prevent Neutrophil-Driven Inflammatory Damage. Cell 177, 541–555.e17 (2019).

30. Lämmermann, T. In the eye of the neutrophil swarm—navigation signals that bring neutrophils together in inflamed and infected tissues. J. Leukoc. Biol. 100, 55–63 (2016).

31. Netea, M. G. et al. Defining trained immunity and its role in health and disease. Nat. Rev. Immunol. 20, 375–388 (2020).

32. Kleinnijenhuis, J. et al. Bacille Calmette-Guérin induces NOD2-dependent nonspecific protection from reinfection via epigenetic reprogramming of monocytes. Proc. Natl. Acad. Sci. **109**, 17537–17542 (2012).

33. Quintin, J., et al. Candida albicans Infection Affords Protection against Reinfection via Functional Reprogramming of Monocytes. Cell Host Microbe 12, 223–232 (2012).

34. Kalafati, L. et al. Innate Immune Training of Granulopoiesis Promotes Anti-tumor Activity. Cell 183, 771–785.e12 (2020).

35. Kato, Y. & Kumanogoh, A. The immune memory of innate immune systems. Int. Immunol. 37, 195–202 (2025).

36. Mitroulis, I. et al. Modulation of Myelopoiesis Progenitors Is an Integral Component of Trained Immunity. Cell 172, 147–161.e12 (2018).

37. Kaufmann, E. et al. BCG Educates Hematopoietic Stem Cells to Generate Protective Innate Immunity against Tuberculosis. Cell 172, 176–190.e19 (2018).

38. Schlüter, T., Van Elsas, Y., Priem, B., Ziogas, A. & Netea, M. G. Trained immunity: induction of an inflammatory memory in disease. Cell Res. 10.1038/s41422-025-01171-y (2025) doi:10.1038/s41422-025-01171-y.

39. Fanucchi, S., Domínguez-Andrés, J., Joosten, L. A. B., Netea, M. G. & Mhlanga, M. M. The Intersection of Epigenetics and Metabolism in Trained Immunity. Immunity 54, 32–43 (2021).

40. Darroch, H. et al. Infection-experienced HSPCs protect against infections by generating neutrophils with enhanced mitochondrial bactericidal activity. *Sci. Adv.* **G**, eadf9904 (2023).

41. Mariotti, B. et al. Innate immune reprogramming in circulating neutrophils of COPD patients. J. Allergy Clin. Immunol. 156, 373–384 (2025).

42. Gomes, M. C., Brokatzky, D., Bielecka, M. K., Wardle, F. C. & Mostowy, S. *Shigella* induces epigenetic reprogramming of zebrafish neutrophils. *Sci. Adv.* 9, eadf9706 (2023).

43. Patel, K. K., Tariveranmoshabad, M., Kadu, S., Shobaki, N. & June, C. From concept to cure: The evolution of CAR-T cell therapy. Mol. Ther. 33, 2123–2140 (2025).

44. Abdin, S. M., Paasch, D. & Lachmann, N. CAR macrophages on a fast track to solid tumor therapy. Nat. Immunol. 25, 11–12 (2024).

45. Chang, Y. et al. CAR-neutrophil mediated delivery of tumor-microenvironment responsive nanodrugs for glioblastoma chemo-immunotherapy. Nat. Commun. 14, 2266 (2023).

46. Chang, Y. et al. Engineering chimeric antigen receptor neutrophils from human pluripotent stem cells for targeted cancer immunotherapy. Cell Rep. 40, 111128 (2022).

47. Majumder, A. et al. iPSC-Derived CAR Neutrophils Possess Potent Activity Against Solid Tumors In Vivo. Blood 144, 86–86 (2024).

48. Sathe, N. et al. Pseudomonas aeruginosa: Infections and novel approaches to treatment “Knowing the enemy” the threat of Pseudomonas aeruginosa and exploring novel approaches to treatment. Infect. Med. 2, 178–194 (2023).

49. Ruffin, M. & Brochiero, E. Repair Process Impairment by Pseudomonas aeruginosa in Epithelial Tissues: Major Features and Potential Therapeutic Avenues. Front. Cell. Infect. Microbiol. 9, 182 (2019).

50. Clatworthy, A. E., et al. *Pseudomonas aeruginosa* Infection of Zebrafish Involves both Host and Pathogen Determinants. Infect. Immun. 77, 1293–1303 (2009).

51. Potvin-Trottier, L., Lord, N. D., Vinnicombe, G. & Paulsson, J. Synchronous long-term oscillations in a synthetic gene circuit. Nature 538, 514–517 (2016).

52. Devanga Ragupathi, N. K., Muthuirulandi Sethuvel, D. P., Inbanathan, F. Y. C Veeraraghavan, B. Accurate differentiation of Escherichia coli and Shigella serogroups: challenges and strategies. New Microbes New Infect. 21, 58–62 (2018).

53. Malet-Engra, G. et al. Collective Cell Motility Promotes Chemotactic Prowess and Resistance to Chemorepulsion. Curr. Biol. 25, 242–250 (2015).

54. Murphy, R. C. & Gijón, M. A. Biosynthesis and metabolism of leukotrienes. Biochem. J. 405, 379–395 (2007).

55. Huang, L. et al. Engineered probiotic Escherichia coli elicits immediate and long- term protection against influenza A virus in mice. Nat. Commun. 15, 6802 (2024).

56. Tamás, S. X. et al. A genetically encoded sensor for visualizing leukotriene B4 gradients in vivo. Nat. Commun. 14, 4610 (2023).

57. Wang, X. et al. Endotoxin-induced autocrine ATP signaling inhibits neutrophil chemotaxis through enhancing myosin light chain phosphorylation. Proc. Natl. Acad. Sci. 114, 4483–4488 (2017).

58. Wang, W. et al. Breakthrough of solid tumor treatment: CAR-NK immunotherapy. Cell Death Discov. 10, 40 (2024).

59. Sterner, R. C. & Sterner, R. M. CAR-T cell therapy: current limitations and potential strategies. Blood Cancer J. 11, 69 (2021).

60. Bradford, Y. M. et al. Zebrafish information network, the knowledgebase for *Danio rerio* research. Genetics 220, iyac016 (2022).

61. Renshaw, S. A. et al. A transgenic zebrafish model of neutrophilic inflammation. Blood 108, 3976–3978 (2006).

62. Bakshi, S. et al. Tracking bacterial lineages in complex and dynamic environments with applications for growth control and persistence. Nat. Microbiol. 6, 783–791 (2021).

63. Sarris, M. et al. Inflammatory Chemokines Direct and Restrict Leukocyte Migration within Live Tissues as Glycan-Bound Gradients. Curr. Biol. 22, 2375–2382 (2012).

64. Livak, K. J. & Schmittgen, T. D. Analysis of Relative Gene Expression Data Using Real- Time Quantitative PCR and the 2−ΔΔCT Method. Methods 25, 402–408 (2001).

65. Williantarra, I., Georgantzoglou, A. & Sarris, M. Visualising Neutrophil Actin Dynamics in Zebrafish in Response to Laser Wounding Using Two-Photon Microscopy. BIO-Protoc. 14, (2024).

66. Paliæ, D., Andreasen, C. B., Ostojić, J., Tell, R. M. & Roth, J. A. Zebrafish (Danio rerio) whole kidney assays to measure neutrophil extracellular trap release and degranulation of primary granules. J. Immunol. Methods **319**, 87–97 (2007).

67. Babatunde, K. A. et al. Chemotaxis and swarming in differentiated HL-60 neutrophil- like cells. Sci. Rep. 11, 778 (2021).

