## Supplementary Material for "In form for a swarm: programmable neutrophil swarming impacts infection outcome"

**Affiliations:**

§ These authors contributed equally

### Supplementary Figures

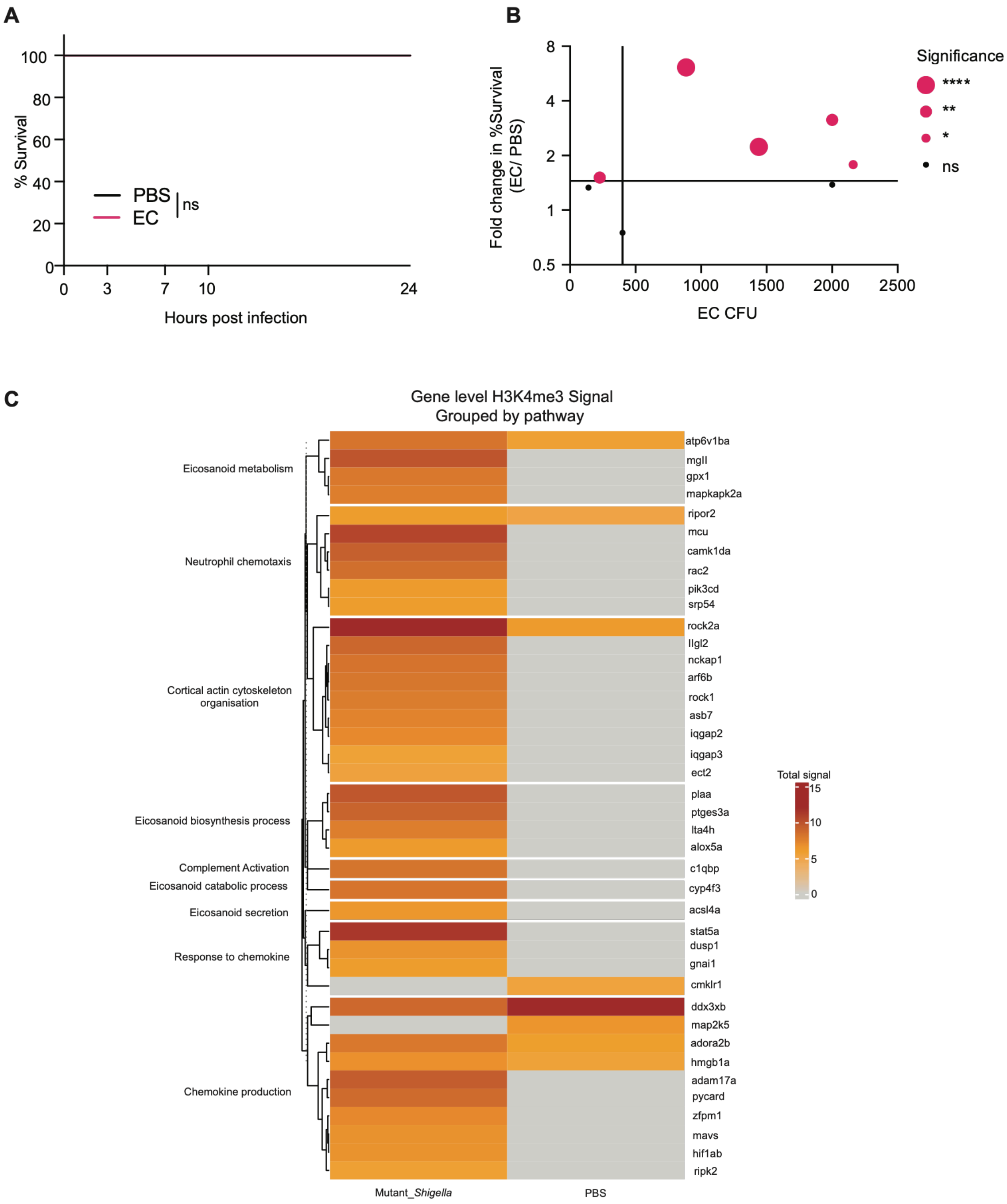

**Figure S1. Phenotypic variations induced by training challenge.**

A. Survival curves from control PBS pre-challenged or trained, *E. Coli* pre-challenged (EC) zebrafish larvae upon PAO1 infection without mechanical wounding. Representative of three independent experiments with $n \geq 7$ larvae per group for each experiment. *Log-rank (Mantel-Cox) test. ns* $p > 0.05$.

B. Relationship between survival advantages acquired by zebrafish larvae upon training and the dosage of *E. coli* injection. Each data point represents an individual training experiment with survival comparisons as in Figure 2B, but with varying dose of *E. coli* for pre-challenge. The degree of statistical significance in survival for PBS pre-challenged versus *E. coli* pre-challenged is shown. *Log-rank (Mantel-Cox) test;* ****$p < 0.0001$, **$p < 0.01$, *$p < 0.05$, ns $p > 0.05$.

C. Heatmap of H3K4me3 ChIP-seq signal intensity at the promoters (≤1 kb) of neutrophil swarming-associated genes (*cyp4f3, lta4h, alox5a, pla2g4aa, alox5ap*). Data were re-analysed from GSE217063 (Gomes et al.), representing neutrophils isolated from zebrafish larvae 48 hours post-injection with either PBS (control) or *Shigella* T3SS-deficient mutant (*mxiD*). Color scale indicates fold increase in signal intensity. Asterisks (*) denote statistically significant enrichment (q-value < 0.05). "Undetected" indicates no peak was called in that region.

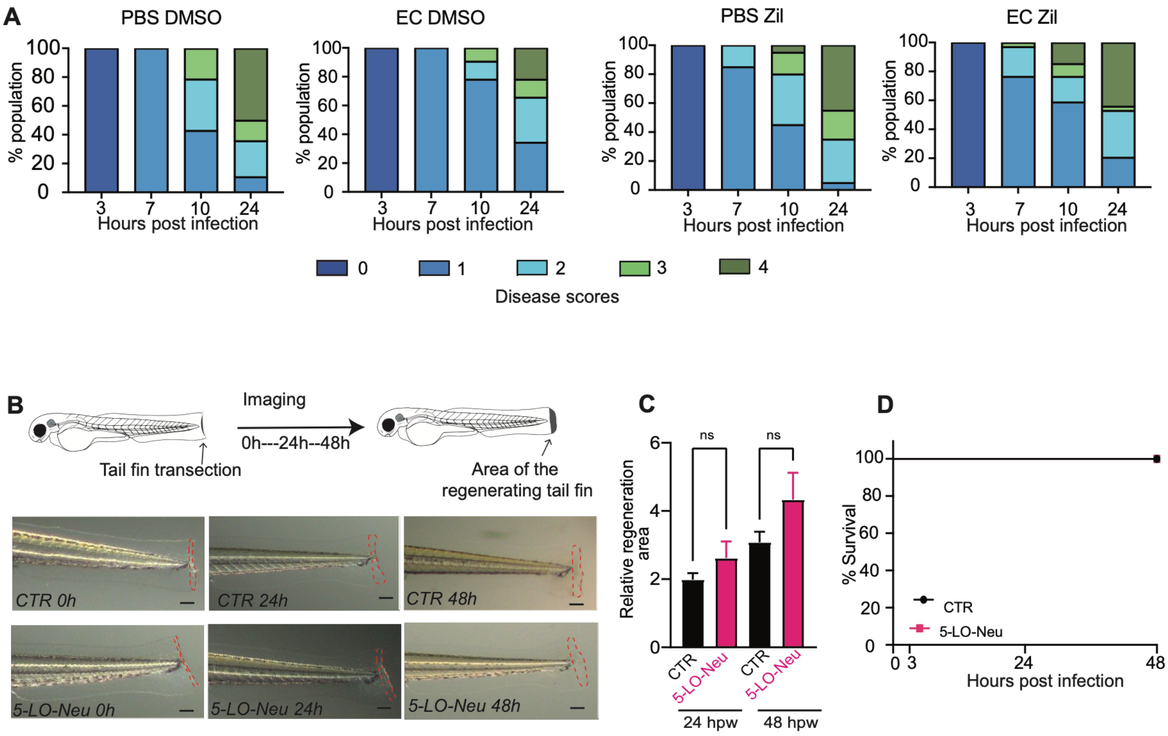


**Figure S2. Disease phenotypes after training challenge and 5-LO engineering.**

A. Disease progression in the control and trained zebrafish larvae at the indicated time points. Representative of three independent experiments with n ≥28 larvae per group for each experiment in J, and n ≥ 20 larvae in K.

B. Tail transection assay to determine area of regeneration in CTR vs. 5-LO-Neu larvae. Schematic depicting the experiment pipeline *(top)*; representative images of CTR and 5-LO-Neu larvae after tail transection (0 h), 24 h post transection and 48 h post transection. Red dashed line indicates the area posterior to the point of transection that was measured as the regeneration area. Scale bar: 100 μm

C. Comparison of the relative regeneration area of the tail fin in CTR and 5-LO-Neu larvae at the indicated time points after tail fin transection. Representative of three independent experiments with n ≥ 24 larvae per group for each experiment. *One-way ANOVA with Tukey's multiple comparisons test, ns: not significant (p>0.05).*

D. Survival curves from CTR and 5-LO-Neu zebrafish larvae upon *P. aeruginosa* infection without mechanical wounding (MW). Representative of three independent experiments with n ≥ 7 larvae per group in each experiment. *Log-rank (Mantel-Cox) test. ns p > 0.05.*

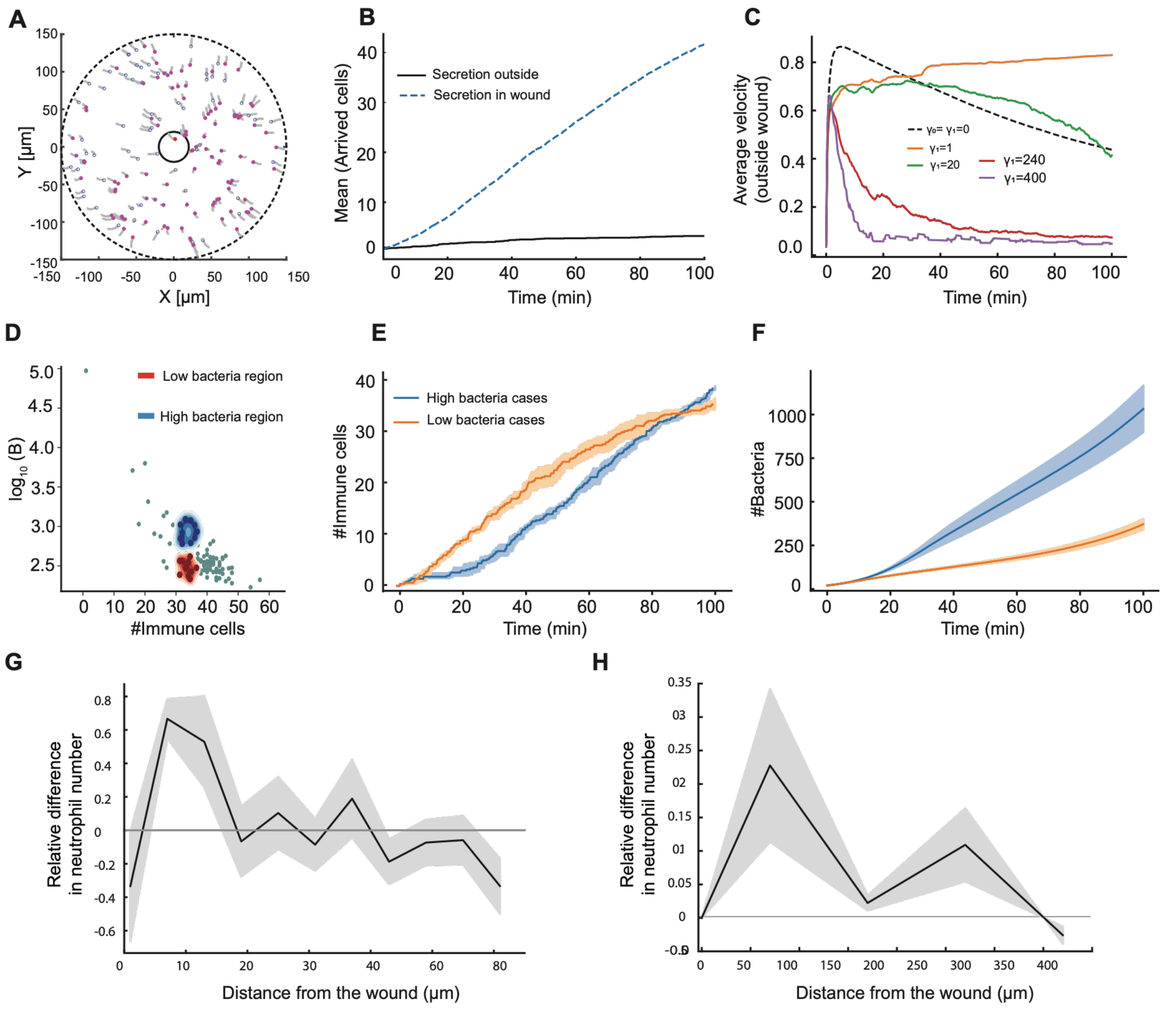


**Figure S3. Modelling the relationship between the dynamics of swarming and bacterial clearance.**

A. Snapshot after 200min of a simulation where the cells are secreting chemoattractant outside the wound. The parameters used here: $r_{max} = 150\mu m$, $r_w = 20\mu m$, $R = 5\mu m$, $c_{wound} = 0.1$, $r_0 = 35\mu m$, $c_{cell} = 35$, $k_0 = 0.055 min^{-1}$ and $v_0 = 15\mu m/min$.

B. Mean number of immune cells arriving at the wound as function of time, for secretion outside the wound (black line) and wound-only secretion (dashed line, with α =35 and parameters mentioned in Table I).

C. The speed towards the wound as function of time, for cells outside the wound, for different values of the stop-signal rate $\gamma_1$ (Eq.5).

D. Simulations that ended (at 100min) in either high (blue region) or low (red region) bacterial burden while having similar final number of recruited immune cells.

E,F. Mean number of recruited immune cells at the wound as function of time, and the corresponding bacterial burden, for the two groups of events in (D).

G. The relative difference in the number of immune cells as function of distance from the wound at the initial time, between the larvae that ended up with low bacteria over those that ended up with high bacteria events of (D).

H. As in (F) from representative experimental examples of the 5-LO engineered neutrophils.

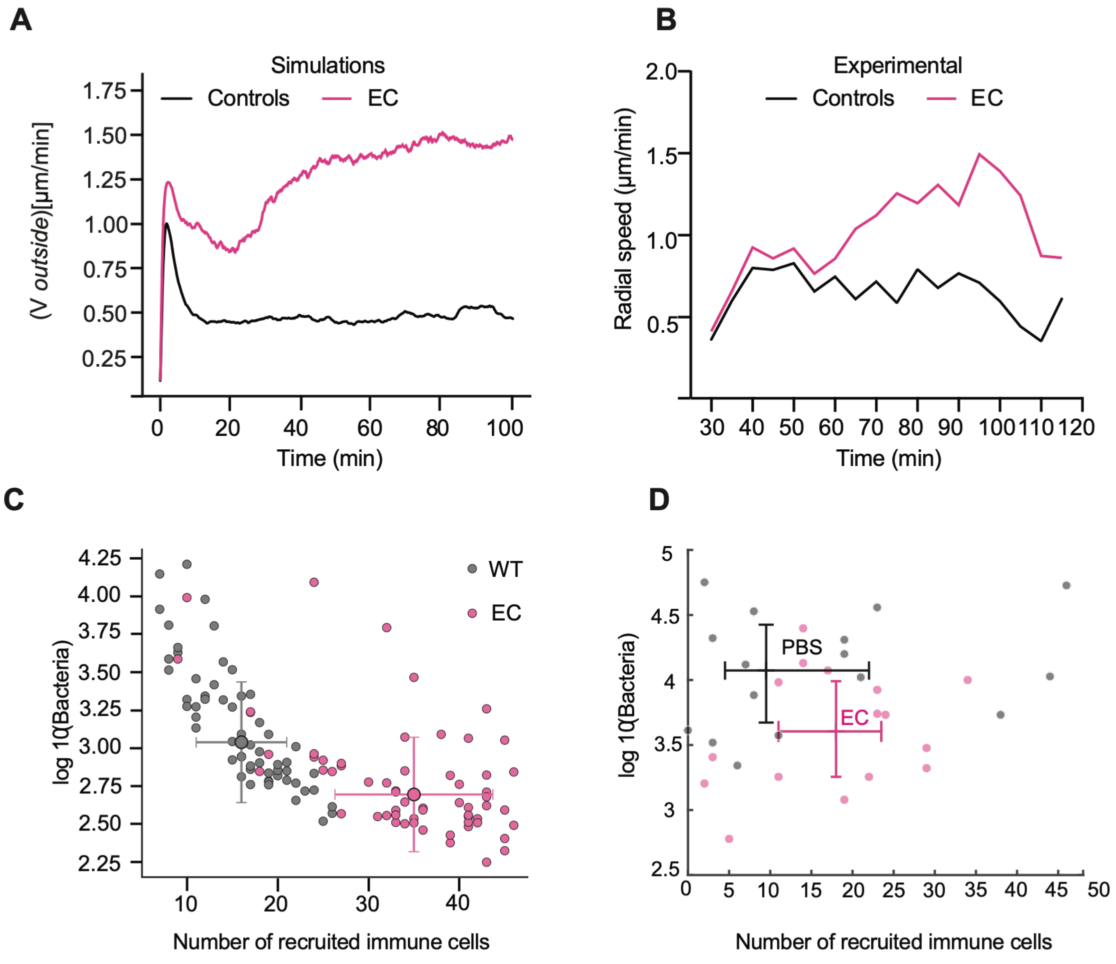


**Figure S4. Modelling complex kinetics of swarming in trained animals.**

A) Simulation results for the mean speed of the cells outside the wound towards the wound, for the WT-like (black) and E.Coli-like (pink) cells. For the E.Coli-like cells the parameters are: $\alpha_0 = 11$, $\alpha_1 = 89$ with $B_s$ =350 (Eq.11).

B) Experimental observations of the mean speed of the cells outside the wound towards the wound, for the WT (black) and E.Coli-treated (pink) fish.

C) Plot of the log of the number of bacteria vs the number of recruited neutrophils (at time=100min) in simulations, for 40 simulations of different random initial conditions. The scatter plot shows the results for WT-like (black) and E.Coli-like (pink) cells. The crosses denote the average values with error bars for standard deviation.

D) As in (C) from the experiments (recruited neutrophils at 120min, and bacteria count at 24hrs). The crosses denote the average values with error bars for standard error of the mean.

#### Supplementary Information on Theory

Additional details of the model simulations

We performed numerical simulations to model the swarming behavior of immune cells toward a wound in the presence of influence of a chemoattractant gradient. The simulation is performed in a two-dimensional circular domain representing the tissue section with an outer radius of $r_{max}$ = 150 μm. The wound region is represented here with a central radius circle $r_w$= 20 μm (as shown in the schematic picture of Fig.5a in the main text), from where the chemoattractant is secreted and diffuses outwards. The concentration profile of the chemoattractant is given by an exponential form (Eq.1 in the Materials and Methods): $c(r) = c_0 \exp[-r/r_0]$. In our simulation, we consider the baseline chemoattractant, $c_0$ = 0.1, with a characteristic decay length of $r_0$ = 35 μm. All the model parameters are summarized in Table I below.

Each immune cell is modeled as a particle of effective radius R = 5 μm. The initial setup begins with a total number (N) = 120 cells randomly distributed in the region between $r_w$ and $r_{max}$. The placement follows a uniform distribution within each annular region, and the angular positions are randomly sampled from $[0, 2\pi)$ to avoid any clustering. The model describes the cell migration by a chemotactic velocity function (Eq.2), $\vec{v}(r) = v_0\,[s(r+R) - s(r-R)]\hat{n}$, as discussed in detail in the Materials and Methods. The local concentration of chemoattractant is governed by a Hill response of first order (Eq.1): $s(r) = c(r) / [c(r) + cs]$. In this case, the maximum motility is $v_0$ = 15 μm/min, and the half-saturation constant of the Hill function is $c_s$ = 1. The strength of chemoattractant secretion per immune cell is controlled by the parameter $\alpha$ (Eq.3).

The simulation consisted of 2,000 steps with a timestep of $\Delta t$ = 0.05 min, for an overall duration of 100 mins. At every timestep, the positions $r_i$ of the cells were computed according to their instantaneous velocity (in μm/min). To prevent cells getting depleted at the outer boundary due to their migration towards the wound, we implemented an open boundary condition through stochastic replenishment: Whenever the density of cells in the outermost annulus dropped below its initial value, new cells were inserted uniformly within that annulus. This approach maintains a quasi-constant density of cells at the periphery while allowing a continuous influx of cells. The wound site ($r \le r_w$) is treated as an absorbing boundary condition. We measure the total number of cells at the wound site by counting the total number of cells that arrived within the wound at a given time.

When immune cells arrive at the wound, they are assumed to immediately begin secreting chemoattractant, as well as switch to stop-signal or attacking states (in the presence of bacteria).
Secretion Outside Wound:
In Fig.S5(A) we demonstrate the dynamics when we assume that the neutrophils can secrete the chemoattractant outside the wound. In this case each cell can switch from non-secreting to secreting depending on the local concentration of the chemoattractant, in the following way

$$k_{secrete}(t) = k_0 \cdot \frac{c(t)}{c(t)+c_s}$$

Each secreting cell contributes to the global concentration field of the chemoattractant, with each cell introducing an exponential profile of concentration around it. The cell's chemotactic migration speed is now not simply directed towards the wound but is aligned towards the local direction of the concentration gradient, which results in the observed clustering of these cells, which prevents their efficient arrival in the wound (Fig.S5(A)).

In this case, the numerical simulation is performed with a similar circular domain with an outer radius = 150 μm and wound radius = 20 μm. For this simulation, 160 cells are initially randomly distributed between $r_w$ and $r_{max}$. Similarly, to maintain the cells density near the outer radius, an open boundary condition is used. New cells are inserted randomly to that outer annulus region whenever the cells density drops below its initial density.

As the immune cells are allowed to secrete chemoattractant outside the wound, thus the resulting chemoattractant concentration can be expressed as the sum of the contribution from the cells that are actively secreting and the contribution from the wound source. The concentration profile of the chemoattractant felt by a cell at radius $r$ thus can be written as,

$c(r,t) = c_{wound}e^{-r/r_0} + \sum_i c_{cell}e^{-|r-r_i|/r_0}$ where there is a constant amplitude source at the wound $c_{wound}$ = 0.1, $r_0$ = 35 μm denotes the decay length of the chemoattractant field and $c_{cell}$ = 35 is the secretion strength of the immune cells secreting outside the wound at $r_i$. Finally, the sum is over all the cells that are secreting at time t. The transition from a non-secreting to a secreting state is given by $w_{secrete}(t) = w_0 * c(r_i,t) \,/\, (c(r_i,t) + C_{RS})$, and this switching depends on the maximal switching rate $w_0$, which is set to 0.055 min$^{-1}$ and $C_{RS}$ (switching half-saturation constant) = 0.6 in our simulation. Accordingly, the rate is converted as a switching probability per timestep, $P_{switch} = w_{secrete}(t) * \Delta t$, with Δt = 0.05 min. In the simulation, the chemotactic velocity of each immune cell is computed based on the directional sensing of the chemoattractant concentration along the cell perimeter, where $N_\theta$ sampled around the circular cell rim. The corresponding unit vector at each of these sampling points is defined as, $\hat{n}_k = (cos\theta_k, sin\theta_k)$concentrations is subsequently computed at the positions, $r_i + R\hat{n}_k$µm is the cells' radius. The directional sensing response is modeled using a Hill function of first order (Eq.1) and the velocity of each cell is then numerically computed as the sum of these local responses along the cell rim: $v_i = v_0(1/N_\theta)\Sigma_{k=1}^{N_\theta}\hat{n}_k c(r_i + R\hat{n}_k)/(c(r_i + R\hat{n}_k) + c_s)$, $v_0$µm/min is the maximal speed. At each timestep, the cells' positions are updated using their instantaneous velocity (in µm/min). Finally, the simulations are performed for 200 minutes with timestep Δt = 0.05 min. In Figure S5(A), we present a snapshot of cells clustering along with their trajectories relative to the initial positions.

| **Parameter** | **Value** | **Unit** |
|---|---|---|
| Cell radius, R | 5 | µm |
| Wound radius, $r_w$ | 20 | µm |
| Tissue radius, $r_{max}$ | 150 | µm |
| Baseline chemoattractant, $c_0$ | 0.1 | a.u. |
| Decay length, $r_0$ | 35 | µm |
| Hill half-saturation, $c_s$ | 1 | a.u. |
| Max. velocity scale, $v_0$ | 15 | µm/min |
| Secretion amplitude, α | 10-200 | a.u. |
| Basal stopping rate, $\gamma_0$ | 1.2 | 1/min |
| Modulation factor, $\gamma_1$ | 240 | 1/min |
| Saturation constant, ns,ss | 100 | -- |
| Initial number of cells, N | 120 | cells |
| Time step, Δt | 0.05 | min |
| Total simulated time, $t_{final}$ | 100 | min |
| **Bacteria-related parameters** | | |
| Initial bacterial load, $B_0$ | 20 | bacteria |
| Bacterial activation saturation, $B_{sat}$ | 1 | bacteria |
| Bacterial interaction saturation, $B_{sat,1}$ | 1 | bacteria |
| Baseline conversion rate, $k_{to\_attack,0}$ | 0.1 | 1/min |
| Baseline bacterial growth, $k_{grow,0}$ | 0.1 | 1/min |
| Attack suppression threshold, $N_{att,grow}$ | 2 | cells |
| Attacking-cell decay rate, β | 0.03 | 1/min |
| Killing efficiency, η | 0.04 | 1/(cell·min) |

### Supplementary Legends

**Movie S1 Examples of neutrophil migration in PBS and *E. coli* pre-challenged larvae**

Examples of neutrophil migration in a Tg(*lyz*:GCamp6F) PBS or *E. coli* pre-challenged and PAO1 infected zebrafish larvae pre and post laser wound (white circle). Frame intervals 20 sec and frame rate is 18.5 fps. Scale bar 50 µm. These videos correspond to the still images in **Figure 2 B.** *E. Coli* example 1 (*left*), *E. Coli* example 2 (*right*) PBS example 1 (*left*) and PBS example 2 (*right*).

**Movie S2 Examples of different neutrophil response to laser wound in control or 5-LO-Neu larvae**

Examples of neutrophil migration in a Tg(*lyz*:GCamp6F) or Tg(*lyz*:GCamp6F) x Tg(*lyz*:tRFP-5LO) PAO1 infected zebrafish larvae pre and post laser wound (white circle). Frame intervals 20 sec and frame rate is 18.5 fps. Scale bar 50 µm. These videos correspond to the still images in **Figure 3 D.** 5-LO-Neu example 1 (*left*), 5-LO-Neu example 2 (*right*), Control (CTR) example 1 (*left*) and CTR example 2 (*right*).

**Movie S3. Example simulation of neutrophil swarming in response to autologous attractant secretion at the target wound.**

Example of a simulation of neutrophils (blue circles) migrating to the wound (black central circle, of radius 20 microns), corresponding to the snapshots in Fig.5(B,C). The cells are randomly placed initially, with uniform average density. As they move towards the wound, they are continuously replenished at random locations around the outer rim (dashed black line, of radius 150 microns) to keep the concentration there constant on average. Cells arriving in the wound (become red circles) secrete chemoattractant that affects the swarming of the cells outside the wound.

**Movie S4. Example of neutrophil response to zymosan *in vitro***

Example of in vitro swarming movies of neutrophils collected from adult zebrafish after intraperitoneal injection with PBS or *E. coli*. These videos correspond to the snapshot in Fig 5 (B). Frame interval 13 sec. Frame rate 20 fps. Scale bar 50 µm. PBS injected fish *(left)* and *E. Coli* injected fish *(right).*
